# Structural Insights into Subunit-Dependent Functional Regulation in Epithelial Sodium Channels

**DOI:** 10.1101/2024.05.28.595834

**Authors:** Alexandra Houser, Isabelle Baconguis

## Abstract

Epithelial sodium channels (ENaC) play a crucial role in Na^+^ reabsorption in mammals. To date, four subunits have been identified—α, β, γ, and δ—believed to form different heteromeric complexes. Currently, only the structure of the αβγ complex is known. To understand how these channels form with varying subunit compositions and define the contribution of each subunit to distinct properties, we co-expressed human δ, β, and γ. Using single-particle cryo-electron microscopy, we observed three distinct ENaC complexes. The structures unveil a pattern in which β and γ positions are conserved among the different complexes while the α position in αβγ trimer is occupied by either δ or another β. The presence of δ induces structural rearrangements in the γ subunit explaining the differences in channel activity observed between αβγ and δβγ channels. These structures define the mechanism by which ENaC subunit composition tunes ENaC function.

## Introduction

The regulation of sodium ion (Na^+^) movement and homeostasis within cells is a fundamental biological process. The epithelial sodium channel (ENaC), a member of the diverse Degenerin/ENaC superfamily, plays a pivotal role in Na^+^ reabsorption.^1^ Dysfunction of ENaC is linked to various diseases, particularly hypertension, which impacts a billion people globally and continues to be the primary cause of morbidity and mortality.^2,3^ ENaCs are composed of three homologous subunits, are sensitive to the pore blocker amiloride, and exhibit selectivity for Na^+^ over K^+^.^1,4–8^ Previous studies identified four subunits - α, β, γ, and δ - and demonstrated a preferential heteromeric assembly of ENaC, forming either αβγ or δβγ complexes.^9–11^ The α and δ subunits are most similar in protein sequence with a homology identity of 37%.^11^ Each subunit confers at least five known distinct channel characteristics when combined with β and γ subunits. First, δ-containing ENaC channels are found in both epithelial and non-epithelial tissues, with the highest expression in reproductive organs, the pancreas, and the brain.^11–14^ On the other hand, α-containing ENaCs are expressed in tight epithelial cells in the kidney, lung, and colon.^15–19^ Second, while δβγ and αβγ selectively permit Na^+^ and Li^+^ over K^+^, the I_Li+/_I_Na+_ permeability ratios for each channel are 0.6 and 2.0, respectively.^11,20,21^ Third, the αβγ channel displays higher sensitivity to the blocker amiloride compared to δβγ by an order of magnitude.^11,21^ Fourth, single channel recordings of the δβγ trimer demonstrate a higher opening probability (*P_o_*) than αβγ^22^. As a result, δβγ is largely unaffected by proteases when compared to αβγ which displays increased activity after exposure to proteases.^23–28^ Lastly, a phenomenon known as Na^+^ self-inhibition is significantly diminished in δβγ, whereas in αβγ, this process reduces channel activity.^29,30^

These variations in expression and channel properties underscore the functional differences between α- and δ-containing ENaCs. While extensive research has been conducted on the structure-function relationship of α-containing ENaC since its identification, there is a significant gap in knowledge concerning δ-containing ENaC. Although all four ENaC subunits are expressed in humans, common mammalian model organisms such as rats and mice do not express δ, hampering in-depth understanding of its function in mammals.^31,32^ To deepen our understanding of ENaC function and, more specifically, to elucidate the molecular basis of distinct properties between δβγ and αβγ, we employed single-particle cryo-electron microscopy and complemented it with electrophysiology to obtain a three-dimensional reconstruction of the assembly and to identify key structural elements that result in the distinct characteristics exhibited by these two ENaC proteins.^33^

## Results

### Stabilization of the trimeric δβγ ENaC complex for structure determination

Wild-type human δ, β, and human γ tagged with enhanced green fluorescent protein (eGFP), collectively designated δβγ_WT_, were expressed in human embryonic kidney (HEK) cells. These cells exhibited a clear preference for Na^+^ over Li^+^ by whole-cell patch-clamp electrophysiology, in agreement with prior δβγ research findings (**Figure 1A and Figure S1A**).^11^ The untagged human δβγ complex exhibited comparable behavior when expressed in *Xenopus laevis* oocytes, with currents measured by two-electrode voltage-clamp electrophysiology (TEVC). This suggests that its function remains intact across different expression systems (**Figure S1B**). Additionally, it demonstrates that tagging the γ subunit does not disrupt its overall function. We expanded our functional characterization to compare the response of δβγ to Zn^2+^, a divalent cation known to elicit a biphasic reaction from αβγ.^34,35^ Upon exposure to 100 µM Zn^2+^, we observed a surprising decrease in δβγ-mediated Na^+^ current, which contrasted with the previously characterized increase in Na^+^ current with αβγ at the same Zn^2+^ concentration ^(^**Figure 1C, D**). We recapitulated a biphasic response pattern with αβγ, while δβγ exhibited a monophasic response with an IC_50_ of 63 µM (**Figure S1C, D**). To our knowledge, this experiment marks the first demonstration of the inhibitory effect of Zn^2+^ on δβγ, shedding light on the significance of this cation to ENaC function and underscoring how variations in subunit composition can yield diverse responses to external modulators.

**Figure 1.**
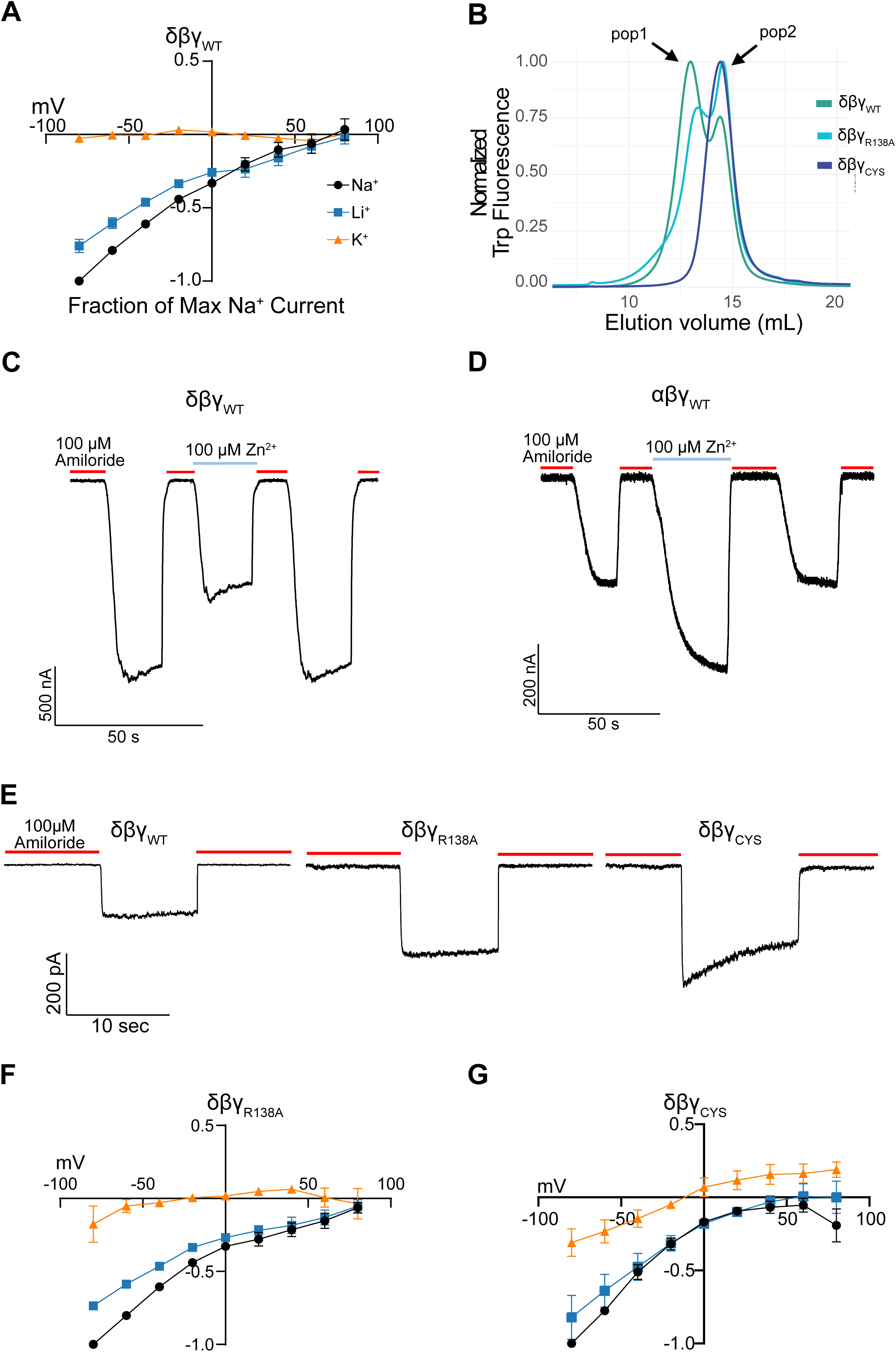
Functional and biochemical characterization of the δβγ complex. **A.** Current-voltage experiment demonstrating that δβγ_WT_ is Na^+^-selective, permeable to Li^+^, and impermeable to K^+^. Voltage potential range used for the experiment is -80 mV to 80 mV in 20 mV increments. The external solutions contained equimolar concentration of Na^+^, Li^+^, and K^+^. Internal solution contained K^+^. Each point is represented as mean ± SEM (n = 7). **B.** Normalized FSEC traces of purified δβγ_WT_, δβγ_R138A_, and δβγ_CYS_monitored on the tryptophan fluorescence channel. Traces were normalized to the height of the peak at 14 mL. **C, D.** Representative traces of δβγ (C) and αβγ (D) with and without 100 μM Zn^2+^. The red and blue lines indicate application of 100 μM amiloride and 100 μM Zn^2+^, respectively. **E.** Representative current traces of δβγ_WT_, δβγ_R138A_, and δβγ_CYS_ expressed in HEK cells and in whole-cell patch-clamp experiments. Holding potential at -60 mV. **F, G.** Current-voltage experiments of δβγ_R138A_ (F) and δβγ_CYS_ (G) using the same conditions in A. Data are represented as mean ± SEM (n = 5 for δβγ_R138A_ and n=5 for δβγ_CYS_). See also Figure S1.

The distinct behavior of the δβγ complex relative to αβγ extends further to its biochemical characteristics.^36,37^ We employed the same purification strategy as the human αβγ, exploiting the high-affinity interaction between a GFP nanobody and the eGFP covalently fused to the γ subunit.^38,39^ When δβγ_WT_ is expressed and purified, two major populations, termed pop1 and pop2, emerge as monitored by fluorescence-detection size-exclusion chromatography (FSEC) (**Figure 1B**).^40^ Pop2 aligns with the expected trimeric ENaC assembly, consistent with the heteromeric αβγ complex. Conversely, pop1 elutes earlier, suggesting a larger complex that is not present in human αβγ expression and purification (**Figure S1E**).^37^ It is important to biochemically and structurally characterize both populations. Consequently, we first focused on strategies to isolate and characterize pop2 because its peak position corresponds to the αβγ trimer, and thus, the predicted position of δβγ trimer.

To improve expression and to generate ideal substrates for detailed structural analysis via single-particle cryo-EM, we systematically designed and screened several other mutant candidates. Numerous investigations have identified and characterized protease recognition sites within the extracellular domain of the γ subunit, one of which is recognized by furin.^23,41–52^ To reduce biochemical variability arising from the putative furin-mediated cleavage during expression, a mutation involving a single residue at the presumed furin site of the γ subunit, specifically R138A, was incorporated producing a construct deemed δβγ_R138A_ (**Figure S1A**). δβγ_R138A_ behaved similarly to δβγ_WT_ biochemically (**Figure 1B**). By replacing cysteines located in the pre-transmembrane domain 1 (pre-TM1) with alanines in all three subunits of δβγ_R138A_, we also created a third candidate, termed δβγ_CYS_ (**Figure 1B**). Biochemical analysis unveiled distinct behavior between the three δβγ constructs based on the distribution between pop1 and pop2 (**Figure 1B**). Despite attempts to exclusively collect pop2 with δβγ_WT_and δβγ_R138A_, there was still contamination from pop1. Successful separation was achieved exclusively with δβγ_CYS_.

Functional characterization of δβγ_R138A_ and δβγ_CYS_ revealed that they exhibited similar amiloride-sensitive inward currents to δβγ_WT_ when expressed in HEK cells (**Figure 1E-G**). However, δβγ_CYS_ behaved differently from δβγ_WT_, displaying a small decay in Na^+^ inward current over time compared to sustained currents observed in δβγ_WT_and δβγ_R138A_ (**Figure 1E**). It has been shown that alterations to the cysteines in the preTM1 region result in channels that favor the closed state in αβγ ENaC.^53,54^ The current decay observed in δβγ_CYS_ aligns with that observation. All three δβγ constructs displayed similar preferences for Na^+^ over Li^+^ and low permeability of K^+^ currents based on bi-ionic experiments (**Figure 1A, F, and G**). Although the three δβγ constructs exhibited similar functional characteristics, the effective separation of pop1 from pop2 when expressing and purifying δβγ_CYS_ prompted us to focus on δβγ_CYS_ for in-depth structural analysis (**Figure 1B**).

### Mapping the ENaC subunits in different heteromeric complexes

We utilized the 10d4 fragment-antigen binding domain (Fab) to label the β subunit and to facilitate particle alignment during cryo-EM data processing.^36^ Reference-free two-dimensional (2D) classification and three-dimensional (3D) reconstruction revealed two different trimeric populations: one with one Fab and the other with two Fabs bound (**Figure S2A**). The trimer with two Fabs bound contains two β, which was an unexpected finding. Beyond just relying on Fab binding, we employed additional strategies to differentiate between the three subunits. The quality of both maps provided sufficient detail to construct the main chain and guide placement of large aromatic residues. Along with the distinct glycosylation pattern of each subunit, this allowed us to confidently determine which subunits belonged to specific positions on the map (**Figure 2A, 2B, and S2B**). After careful inspection of these maps, we determined that the monoFab trimer comprises δ, β, and γ, which we refer to as δβγ_CYS_. The diFab complex consists of two β and a γ, and we call this trimer ββγ_CYS_ (**Figure 2A-D and S2A**).

**Figure 2.**
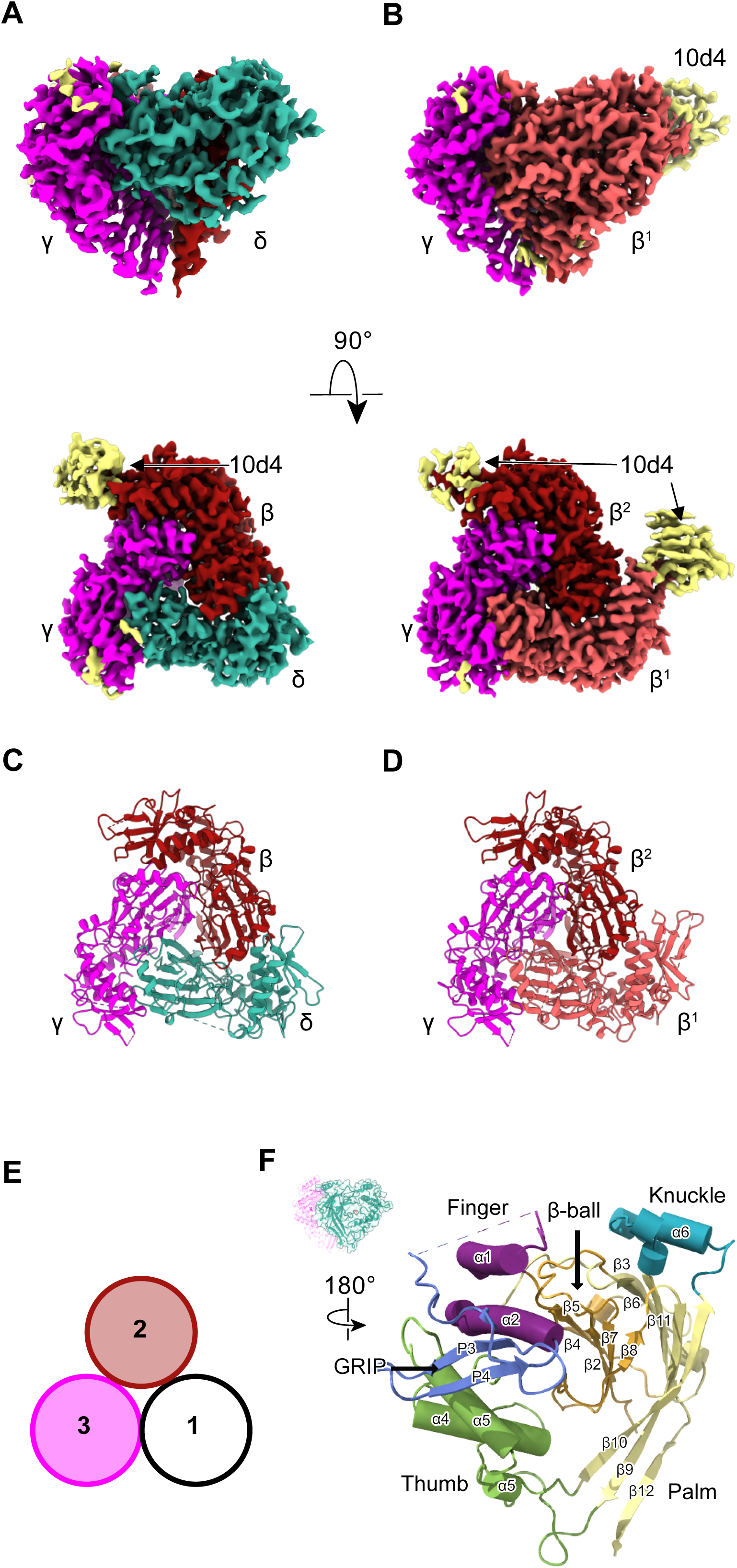
Cryo-EM analysis reveal two different heteromeric complexes. **A**. Cryo-EM map of the extracellular domain of δβγ_CYS_ viewed parallel and perpendicular to the membrane plane viewed from the extracellular side, and in surface representation. The subunits δ, β, and γ are colored teal, red, and magenta, respectively. The 10d4 Fab bound to the β subunit is colored yellow. **B**. Cryo-EM map of ββγ_CYS_ viewed similarly as in A. The β^1^ subunit is colored salmon. Only a small segment of the 10d4 Fab, colored yellow, is resolved after local refinement. **C, D**. The models of the extracellular domains of δβγ_CYS_ (A) and ββγ_CYS_ (B) viewed from the extracellular side and down the pore axis. They are in cartoon representation and colored as in Figure 2A and B **E**. Schematic illustration viewed from the extracellular side and looking down the pore axis of the three positions of the subunits in ENaC denoted as positions 1, 2, and 3. **F**. Inset: An overall view of the δβγ_CYS_ extracellular domain in cartoon representation. The close-up view of δ is rotated 180° relative to the view in the inset. The δ subunit follows a similar architecture as other ENaC subunits consisting of domains arranged like a hand grasping a ball. The unique domain first characterized in the human αβγ structure, the GRIP domain colored blue, is conserved in δ. The knuckle, finger, thumb, palm, and β-ball are colored cyan, purple, green, yellow, and orange, respectively. See also Figure S2.

The majority of the extracellular domains of δβγ_CYS_ and ββγ_CYS_ were resolved with reported resolutions of 3.4 Å and 3.1 Å, respectively, based on the gold standard Fourier shell correlation (GSFSC) at 0.143 (**Figure 2A, B, S2C-H, and Table S1**). However, like the previous human αβγ cryo-EM studies, the transmembrane domains remain unresolved.^36,37^ Based on the constructed model, the arrangement of the δβγ_CYS_ subunit is consistent with the αβγ arrangement, with the δ-β-γ orientation following a counterclockwise direction when viewed from the extracellular side and down the pore axis (**Figure 2C**).^36^ In comparison to the αβγ arrangement, the δ subunit occupies a position equivalent to that of α, while β and γ subunits maintain the same positions (**Figure 2A-D**).

The second trimer we observed in our dataset, ββγ_CYS_, offers additional insight into how compositional differences in subunit stoichiometries can give rise to structural differences. We did not observe a trimeric complex comprising one β and two γ subunits or a complex consisting of three γ subunits (**Figure S2A**). The presence of 10d4 bound to the β subunit in all trimers indicated that the binding of the Fab is not dependent on the subunit composition of the trimer. To simplify the analysis, we designated the position occupied by α, δ, and β as position 1, and the positions consistently occupied by β and γ subunits as positions 2 and 3, respectively **(Figure 2E**). To distinguish between the two β subunits within the ββγ_CYS_ complex, we denote the β subunit in position 1 as β^1^ and the one in position 2 as β^2^ (**Figure 2B, D**).

### Architecture of the **δ** subunit

The δ subunit shares a structural resemblance with α, β, and γ, exhibiting subunit domains arranged in a hand-like configuration clenching a ball replete with β strands, first observed in the crystal structure of chicken ASIC1 (**Figure 2F**).^4^ Similar to α, β, and γ, δ also comprises extended anti-parallel helices, namely α1 and α2, which form part of the finger domain. These helices resemble a wall positioned between the functionally important β6-β7 loop and the thumb domain.^36^ Like β, δ is thought to be insensitive to proteases but harbors the **g**ating **r**elief of **i**nhibition by **p**roteolysis domain, or GRIP, which forms extensive contacts with the finger and thumb domains.^36^ The knuckle sits atop the expansive palm domain and makes contacts with the adjacent γ finger (**Figure 2F)**. While the upper palm domain is well-resolved, the lower palm domain in δ is disordered, hindering precise positioning of the β strands that directly connect to the transmembrane domains and wrist region (**Figure 2A**). This stands in stark contrast to β and γ, which exhibit clear β strands in the lower palm domains. Similarly, the αβγ maps also displayed defined features in the lower palm domain.^36,37^ The disorder observed in the lower palm domain in δ may suggest that this region adopts more conformations compared to the other subunits. This unique characteristic of δ likely contributes to the distinct properties of δβγ compared to αβγ.^9–28^

### Position 1 subunit modulates conformational changes

To compare the trimers, we leveraged the well-resolved map regions encompassing the α2 helices within the finger domains of each subunit. Our rationale was grounded in the superior map quality in this specific region, allowing us to assign and position residues, particularly the bulky aromatic side chains (**Figure S3**). With this approach, we focused on evaluating the distances of the Cα atoms of conserved tryptophans between associated subunits and then compared the relative distance between each position in the three trimers. We chose these key tryptophans—Trp232 (δ), Trp251 (α), Trp218 (β), and Trp229 (γ)—because they are located in a region where domains believed to regulate ENaC activity converge, incorporating the thumb, finger, and GRIP domains.^36,37^ Collectively, these conserved tryptophans form a triangle, which we refer to as TriTrp. The TriTrp triangle exhibited distinct side lengths between the trimers, with the δβγ_CYS_ TriTrp triangle featuring longer sides than those of αβγ (PDB: 6WTH) and ββγ_CYS_ (**Figure 3A-C**). The TriTrp triangle within ββγ_CYS_forms an isosceles triangle with two sides measuring about 53 Å (**Figure 3B**). The different TriTrp distances between positions 2 and 3, which are consistently occupied by β and γ, in the three trimers suggest that the subunit in position 1 can alter domain positions in the trimer.

**Figure 3.**
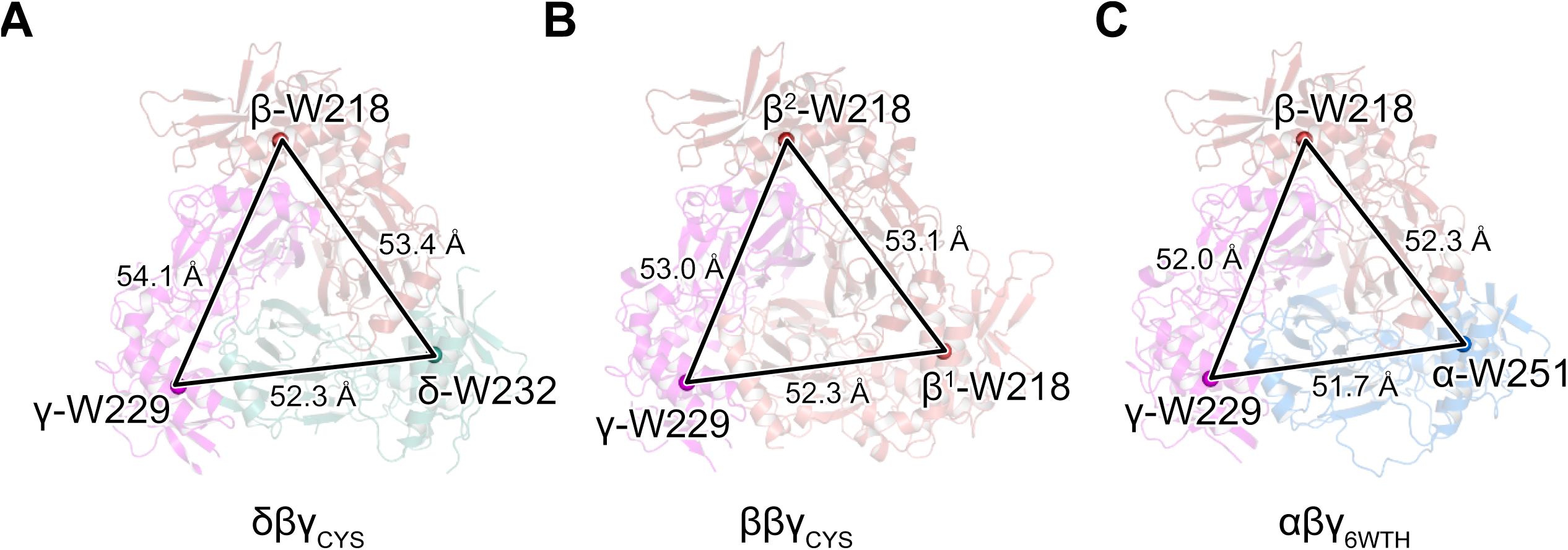
Position 1 subunit mediates changes in the extracellular domain. **A-C.** Comparison of the extracellular domains of δβγCYS (A), ββγCYS (B), and αβγ (C, PDB: 6WTH). The Cα positions of the tryptophans belonging to the TriTrp are shown as spheres. The distances between the Cα atoms are shown in Å. See also Figure S3.

To identify regions contributing to changes in the extracellular domain, we systematically compared the corresponding subunit at positions 1-3 of each trimer by superposing them. We observed conformational differences in the finger domains of the subunits occupying positions 1 and 3 (**Figure 4, Table S2**). Conversely, in position 2, the β subunits exhibited almost identical structures, demonstrating an overall root-mean-square-deviation (RMSD) of less than 0.50 Å (**Figure S4A**). The remarkable similarity of the β subunits in position 2 from the three trimers served two distinct purposes. First, it demonstrated that the β subunit undergoes minimal conformational changes even when the subunit in position 1 changes. Second, it confirmed that the cryo-EM maps and resulting models were consistent and on the same map scale. Thus, observed rigid-body differences between the trimers are valid and not artifacts derived from map scale discrepancies.

**Figure 4.**
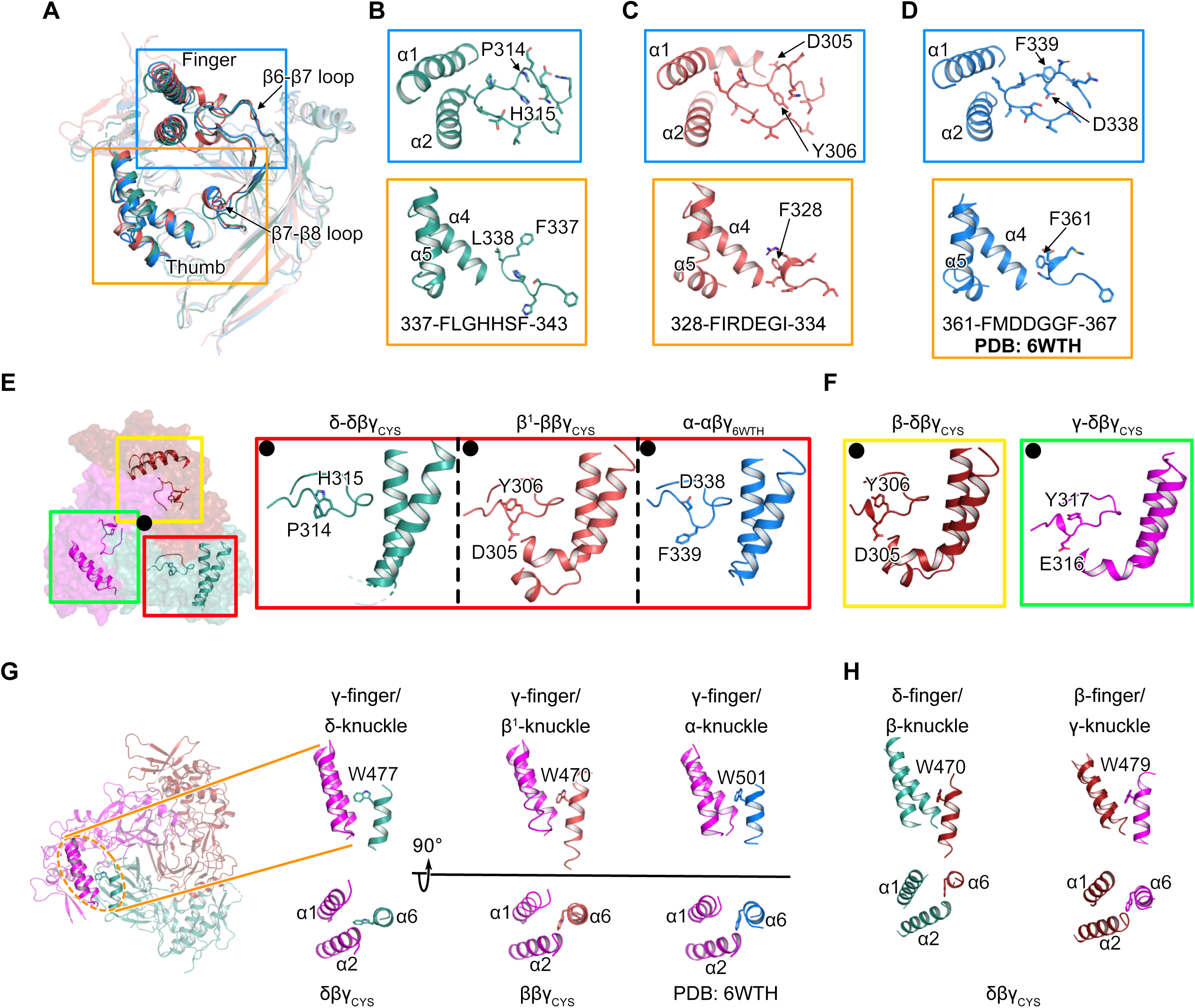
Unique conformational features in δ contribute to changes in the extracellular domains. **A**. Superposition of all position 1 subunits show differences in the finger and β-ball domains. The δ, β^1^, and α are colored teal, salmon, and blue. The subunits are shown in cartoon representation. **B-D**. Close-up view of boxed regions in A belonging to δ (B), β^1^ (C), and α (D). The sidechains of β6-β7 and β7-β8 loop residues are shown in sticks representation. A 7-residue segment of the β7-β8 loop is shown along with the corresponding sequence. **E**. View looking down the pore axis to show relative positions of the finger domains in positions 1-3. The close-up views of the boxed area in position 1 are displayed. The sidechains of residues at corresponding positions in δ, β1, and α are depicted in stick form to illustrate the orientation of their side chains. The black dot marks the relative direction of the pore axis position. **F**. Close-up views of the boxed regions in E of positions 2 and 3 in the δβγ_CYS_ structure. The views are rotated ∼120° clockwise (yellow) or counterclockwise (green) relative to the panels in E highlighted with a red box. **G**. Illustration of the position 1/position 3 interface focusing on the knuckle/finger domain contacts. Only the sidechain of the conserved tryptophan in the knuckle domain is displayed in stick form while the rest of the region is shown in cartoon. **H**. Equivalent interfaces in position 1/position 2 and position 2/position 3 in δβγ_CYS_are shown and represented similarly as in G. See also Figure S4.

Comparing the subunits occupying position 1, α, δ, and β^1^ differ in sequence and molecular interactions contributing to variations in the overall trimer conformations. Evaluating the respective domains revealed significant differences, especially the β-ball domains (**Table S2**). Because of the location of the β-ball, nestled between the structurally homogeneous upper palm domain on one side and the gating domains on the other, we opted to concentrate on this region and explore how its architecture could lead to differences in the conformation of the surrounding domains between the three subunits (**Figure 4A**). Positioned beneath the finger domain, a region that is thought to mediate gating in ENaC and encompasses the β6-β7 loop and the α1 and α2 helices, the β-ball in δ displays a distinct arrangement compared to α and β^1^ (**Figure 4B-D**). Differences in molecular interaction were observed near the interface between the thumb domain and the β7-β8 loops. In this loop, α and β^1^ contain acidic residues while histidines occupy the corresponding positions in δ. With a RMSD of over 3 Å, these loops present distinct conformations and establish varied contacts with the base of the thumb domains. Where phenylalanines of both α and β^1^ (αPhe361 and β^1^Phe328) forge contacts with the base of the α4 helix of the thumb region, δ has a leucine (Leu338) in the equivalent position. The adjacent δPhe337 faces away from the base of α4 and forms contacts with the residues in the β-ball (**Figure 4B**).

The conformation of the finger domain remains similar between δ and α, but there are clear sequence variations likely contributing to the differences in functional behavior of channels containing δ or α. For instance, the β6-β7 loop, crucial for mediating Na^+^-self inhibition in αβγ, features a putative cation site near Asp338 in the structure (**Figure 4D**).^37,55,56^ However, in the δ loop, the corresponding residue is a proline, which lacks a negative charge to directly interact with Na^+^ based on the structure (**Figure 4B**). In this region, δ differs in conformation from α, where δ-His315 points toward the pore axis while α-Phe339 is oriented away (**Figure 4E**). The β6-β7 loop of δ closely resembles that of β^1^, which is akin to those found in position 2 β and γ in all three trimers suggesting that the swapped conformation observed in α Asp338-Phe339 is not unique to the subunit occupying position 1 but more specific to α (**Figure 4E, F, and S4B**).^37^ In β and γ, the section leading up to the extended α1 helix within the finger domain forms a helical structure absent in both δ and α. This section tightly surrounds the β6-β7 loop, likely hindering the accommodation of a large aromatic residue in the β6-β7 loop.

The placement of the β6-β7 loop must affect the position of the α-helical segments of the finger domain.^56^ Comparison of the Cα positions of the core finger domain in the three subunits occupying position 1 demonstrated that β^1^is the most different (**Table S2**, **Figure 4A-D**). This difference observed in β^1^ is likely influenced by the interaction between the β6-β7 loop and the α1 helix, facilitated by the presence of bulkier side chains such as Phe309 and Met119, respectively. In contrast, the equivalent residues in δ and α involve smaller side chains, such as leucine, alanine, and threonine (**Figure 4B-D and S4C**). The conformational differences are more pronounced when shifting away from the central pore axis toward the periphery, where the thumb domain is located. While α and β^1^ exhibit similar conformations, δ displays the most distinct conformation, characterized by a roughly 15° rotation of the α5 helix within the thumb domain (**Figure S4D**).

The differences in the overall placement of the finger domains among the subunits at position 1 do not adequately explain the expansion of the extracellular domain observed in the TriTrp distances of δβγ_CYS_ and ββγ_CYS_. Because the knuckle domains of each subunit establish extensive contacts with the neighboring finger domains, our subsequent analysis focused on understanding how the knuckle domain at position 1 might mediate variations in the domain positions surrounding it. The knuckle domain in position 1 abuts the finger domain of the γ subunit in position 3. We discovered that a conserved tryptophan in the knuckle domain that assumes similar rotamers packs tightly against the α6 helix in α, β, and γ (**Figure 4G, H and S4E**). However, the tryptophan found in δ exhibits a rotamer that extends away from the helix main chain, likely affecting the position1-position3 interface (**Figure 4G and S3A**).

### The **γ** subunit finger domains adopt different conformations

To identify conformational shifts in the subunits at position 3, we aligned the γ subunits, examined the resulting RMSD values for each subunit domain, and found the finger domain exhibited the greatest difference (**Table S2**). Within the finger domain, we identified variations in the positions of three components: the β6-β7 loop, α1, and α2 helices (**Figure 5A**). The β6-β7 loops displayed marked differences, with the main chain involving Val322 and Ser323 adopting swapped positions in γ-δβγ_CYS_ relative to γ-ββγ_CYS_ and γ-αβγ_6WTH_ (**Figure 5B-D, S3C, and S5A**). This alteration influences the position of Phe320, which directly interacts with the α1 and α2 helices of the finger domain. When assessing the angle formed by the two helices within each subunit, we observed that α1 and α2 create a 23° angle in γ-αβγ_6WTH_, while the equivalent helices in γ-δβγ_CYS_ and γ-ββγ_CYS_form 11° and 16° angles (**Figure 5B-D**). Examining this angular change within the context of a trimer revealed that the α2 helices in the γ-δβγ_CYS_and γ-ββγ_CYS_ are pushed outward from the pore axis when compared to that of γ-αβγ_6WTH_ by about 10° and 4°, respectively (**Figure 5E, F**). The substantial rotation of the finger helices stands in stark contrast to the positioning of the thumb domains in γ. We did not observe substantial rotations of the thumb domains across all three γ subunits (**Figure S5B**).

**Figure 5.**
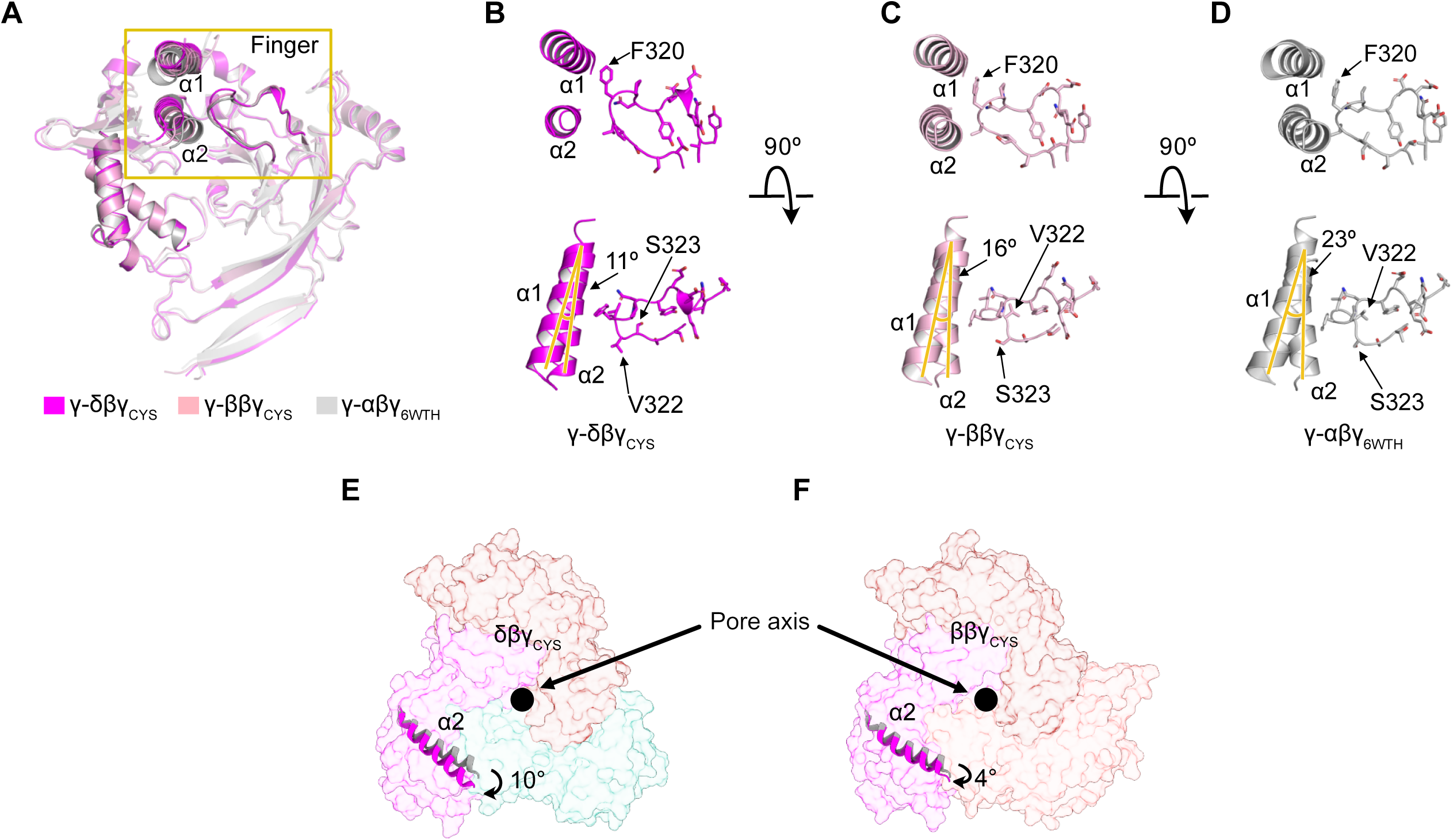
The γ finger domain is altered in the presence of δ. A. Superposition of all γ subunits in position 3 from the three trimers. The subunits are displayed in cartoon representation and colored magenta (δβγ_CYS_), light pink (ββγ_CYS_), and gray (αβγ_6WTH_). B-D. Close-up views of the boxed region in A. The sidechains of residues in β6-β7 loop in the finger domain are shown in stick form to demonstrate the positions of the side chains. E, F. Surface representation of the δβγ_CYS_ (E) and ββγ_CYS_ (F) models is shown. The positions of the γ-α2 helices in δβγ_CYS_ and ββγ_CYS_ relative to those of αβγ_6WTH_ are displayed. See also Figure S5.

Comparison of the maps among the three trimers unveils differences in the GRIP domains of the γ subunit (**Figure S5C**). The γ subunits in the δβγ_CYS_ and ββγ_CYS_ complexes have similar map characteristics in the GRIP domain, featuring an additional element not observed in the αβγ_6WTH_complex. To determine the identity of this map feature, we generated a model of the δβγ trimer using AlphaFold2.^57^ In the predicted model, the γ-GRIP contains additional β strands, with the unmodeled features likely representing one of the predicted short β strands (**Figure S5D**). Among the three trimeric complexes resolved thus far, only αβγ_6WTH_ displays a unique γ-GRIP conformation. The γ-GRIP of αβγ_6WTH_ extends approximately 10 Å further towards the C-terminal end of α2 compared to the γ-GRIP of δβγ_CYS_ and ββγ_CYS_ (**Figure S5C**). While our observation is limited to a portion of the γ-GRIP within the context of the ENaC trimeric assemblies, functional data from prior studies have indicated differences between δβγ and αβγ. Specifically, the γ-GRIP in γ-αβγ exhibits sensitivity to proteases, whereas that in γ-δβγ does not.^28^ Additionally, our study unveils contrasting reactions of δβγ and αβγ to 100 µM Zn^2+^ (**Figure 1C**). Previous investigations into the influence of Zn^2+^ on αβγ activity have highlighted binding sites in the γ finger, GRIP, and palm domains.^34,35^ Specifically, histidines in the γ-GRIP of αβγ were implicated in mediating the stimulatory Zn^2+^ site. These sites are currently not resolved in any of the cryo-EM maps.^36,37^ Nevertheless, the distinct positions of the finger domain of γ and differences in map features in the GRIP segment between δβγ_CYS_ and αβγ_6WTH_ offer insights into how these heteromeric complexes could manifest divergent functional responses to various stimuli.

### The **β** subunit as a structural scaffold

The superposition of the trimers demonstrated positional differences in the gating domains, which are most discernible in position 1 (**Table S2**). We took advantage of the uniform conformation of the β subunits in position 2 and used them as references for aligning the trimers to precisely pinpoint how the subunits in position 1 and 3 rearrange in relation to position 2. This allowed us to effectively identify sites that consist structural variability within both position 1 and 3 (**Table S2**). For clarity and to distinguish between the two ends of the finger domain, we adopted the conventions of (+) and (-). We designated the region of the finger domain making extensive contacts with the thumb as the “+” end and the area interacting with the neighboring knuckle domain as the “-” end (**Figure 6A**). We observed the most significant differences in position 1, especially when comparing the δ and β^1^domains with the α subunit. While δ and β^1^ have differences in helical positions comprising primarily of rotational components, it is when comparing these subunits with α that striking differences are observed (**Figure 6B**). The finger and thumb helices of δ and β^1^—α1, α2, α3, and α4—sit higher than those of α, and exhibit a rotation relative to α, with the “-” end acting as the pivot point (**Figure 6C, D**). Consequently, the “+” end exhibits the greatest difference in position. Interacting with the “+” end, the thumb domain positions of δ and β^1^ are also different compared to α. These distinct domain placements can be summarized by determining the position of the centers of mass of the helical components of the finger and thumb domains from each subunit. The resulting position is located beneath α2. A comparison of these positions shows that the greatest difference is between α and β^1^, with a distance of 5 Å (**Figure 6B-D**). While the “+” region of the finger in δ and β^1^ shows the greatest variation in position, the corresponding areas in position 3 remain stationary (**Figure 6B-D**). Instead, it is the “-” end of the finger domain that demonstrates the largest positional difference, rotating away from the protein core, akin to observations seen when individual subunits are superimposed (**Figure 5**).

**Figure 6.**
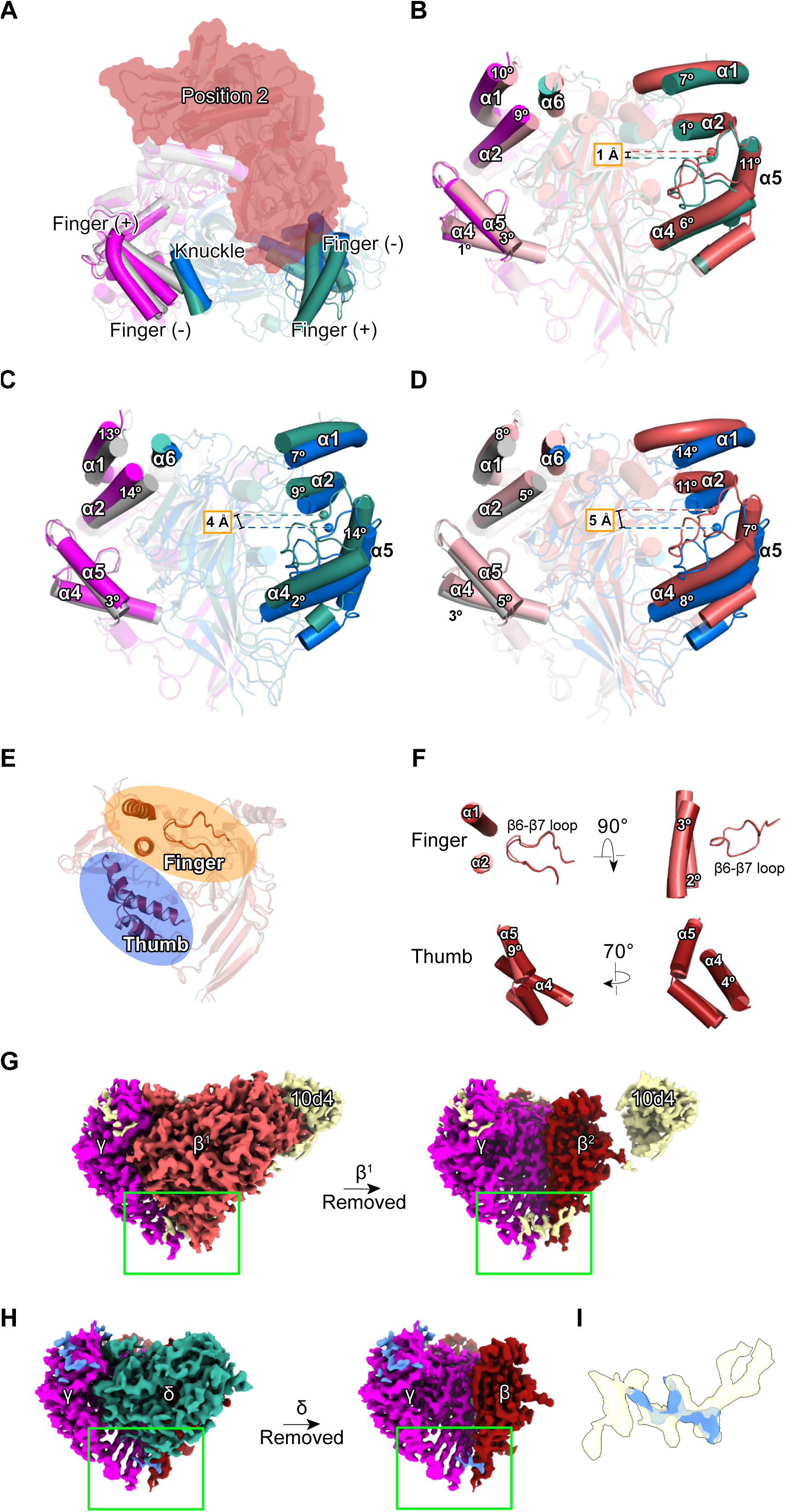
Trimer superpositions using the β subunit in position 2 as reference reveal global rearrangements in position 1. **A.** View looking down the pore axis from the extracellular side. The δβγ_CYS_ and αβγ_6WTH_ trimers are shown in cartoon and subunits are colored as in figures 4 and 5. The helices are shown as cylinders. The β subunit occupying position 2 is colored red. The knuckle, finger, and thumb are opaque while the rest of the extracellular domains are transparent for clarity. **B-D.** Comparison of the finger and thumb domains of position 1 and 3 subunits in δβγ_CYS_ vs ββγ_CYS_ (B), δβγ_CYS_vs αβγ_6WTH_ (C), and ββγ_CYS_ vs αβγ_6WTH_ (D). The β subunit in position 2 is omitted for clarity. The centers of mass of the finger and thumb domain helices, calculated using Pymol, are represented as spheres. The solid bars indicate the distances between the centers of mass. Angle between corresponding helices of the finger and thumb domains are shown. **E.** Overlay of β^1^and β^2^ subunits. **F.** Close-up views of the finger and thumb domains shown in E. **G, H.** Cryo-EM maps of ββγ_CYS_ (G) and δβγ_CYS_ (H) showing a peptide-like map feature in the palm domain that is not observed in αβγ. Position 1 subunits are removed for clarity (right image). The molecules in ββγ_CYS_(G) and δβγ_CYS_ (H) are colored yellow and blue, respectively. **I.** Close-up view and superposition of the features observed in the lower palm domain shown in G and H. See also Figure S6.

Overall, each position within the trimer serves distinct structural roles. The subunit in position 1 influences the overall conformation of the trimer while the position 2 β subunit acts as a structural scaffold. The larger differences of domain positions observed in position 1 likely contribute to the functional differences observed in electrophysiological experiments. Position 3, occupied by the γ subunit, comprises two regions: one anchored by its interaction with the position 2 β subunit, and another adapting to the subunit at position 1 (**Figure 6A**). Position 3 likely serves to regulate or balance the magnitude of influence imposed by the subunits in positions 1 and 2.

The structural rigidity seen in β^2^ subunits is mirrored in the β^1^ subunit, showing a remarkably similar conformation. Although there are minor distinctions between them, the overall resemblance is striking. (**Figure 6E**). While subtle changes were identified in the finger domain, the most observable change is in the α5 thumb domain helix (**Figure 6F**). Here, the middle segment of the helix is rotated by approximately 9°, while its N- and C-termini remain fixed, establishing connections with the α4 and finger domain at the N-terminal end, and the wrist region at the C-terminal end.

We observed small rigid-body rearrangements in the palm domains of β^1^ and β^2^. However, the most striking observation near the lower palm domain is a feature resembling a peptide. This molecule wraps around β^1^ and forms multiple contacts with β^2^ (**Figure 6G, S6A, and S6B**). Due to limitations in map quality, we cannot fully trace this new feature. However, when we examined similar regions in δβγ_CYS_ and αβγ_6WTH_, we found that δβγ_CYS_ contained a smaller but similar feature, while αβγ_6WTH_ showed no equivalent feature near the lower palm domain (**Figure 6H, I**). The presence of this peptide-like feature in both δβγ_CYS_ and ββγ_CYS_ could partly explain the conformational differences observed in these trimers compared to αβγ_6WTH_. The location of this molecule is strategic as it binds in the lower palm domain poised to influence the arrangement of β strands connected to the transmembrane helices. It is unclear whether this molecule is a ligand regulating channel activity or a chaperone-like structure involved in assembly and trafficking. Further research is required to identify the molecule and to characterize its role in ENaC function.

To test whether the ββγ trimer could contribute to the Na^+^ currents we recorded from δβγ_-_expressing HEK293 cells, we infected cells with a construct containing only β and γ (**Figure 1**). We did not detect meaningful amiloride-sensitive currents (**Figure S6C)**. These infected cells behaved similarly to uninfected HEK293 cells (**Figure S6D**). Immunofluorescence light microscopy experiments that leverage our 10d4 antibody aligned with our electrophysiology data demonstrating no surface expression when only β and γ were expressed in HEK cells (**Figure S6E, F**). This confirms that previously described amiloride-sensitive currents were solely a reflection of δβγ activity (**Figure 1A, C, and E-G**). While there has been work in oocytes showing amiloride-sensitive currents with ENaC homotrimers of all subunits and currents with oocytes containing β and γ, it remains to be seen whether ββγ heterotrimers are physiologically relevant^58^.

### The **βγ** dimer provides structural insight into ENaC assembly

Functional and biochemical examinations into ENaC processing have shed light on the specific pathways taken during ENaC protein expression.^52,59–63^ Studies have shown that ENaC subunits are present in both immature and mature forms, distinguished by observed glycosylation patterns.^63,64^ Furthermore, it is also known that despite abundant ENaC expression, only a small fraction reaches the plasma membrane.^65^ However, the precise oligomeric assembly of ENaC when it resides intracellularly remains elusive. We identified a complex within the earlier eluted peak, pop1, which does not assemble as a trimer (**Figure 1B**). Upon closer examination of pop1 from δβγ_WT_ by Cryo-EM, it became evident that its larger size by FSEC compared to the trimers is attributed to the presence of two βγ dimers interacting via the γ subunit, with each β subunit associated with one 10d4 Fab (**Figure 7A**). The dimer-dimer complex lacks a subunit in position 1 (**Figure 7B**). Instead, the space is partially occupied by the γ subunit belonging to the neighboring dimer. The interdimer contact between the two γ subunits is mediated by a segment not observed in any of the trimers—αβγ_6WTH_, δβγ_CYS_, and ββγ_CYS_—likely because this region is no longer stabilized when assembled in a trimer. Close inspection of this region revealed that the features emanate from each γ-GRIP and resemble stretches of β strands. These long β strands from both γ-GRIP interlock stabilizing the dimer-dimer interface (**Figure 7**).

**Figure 7.**
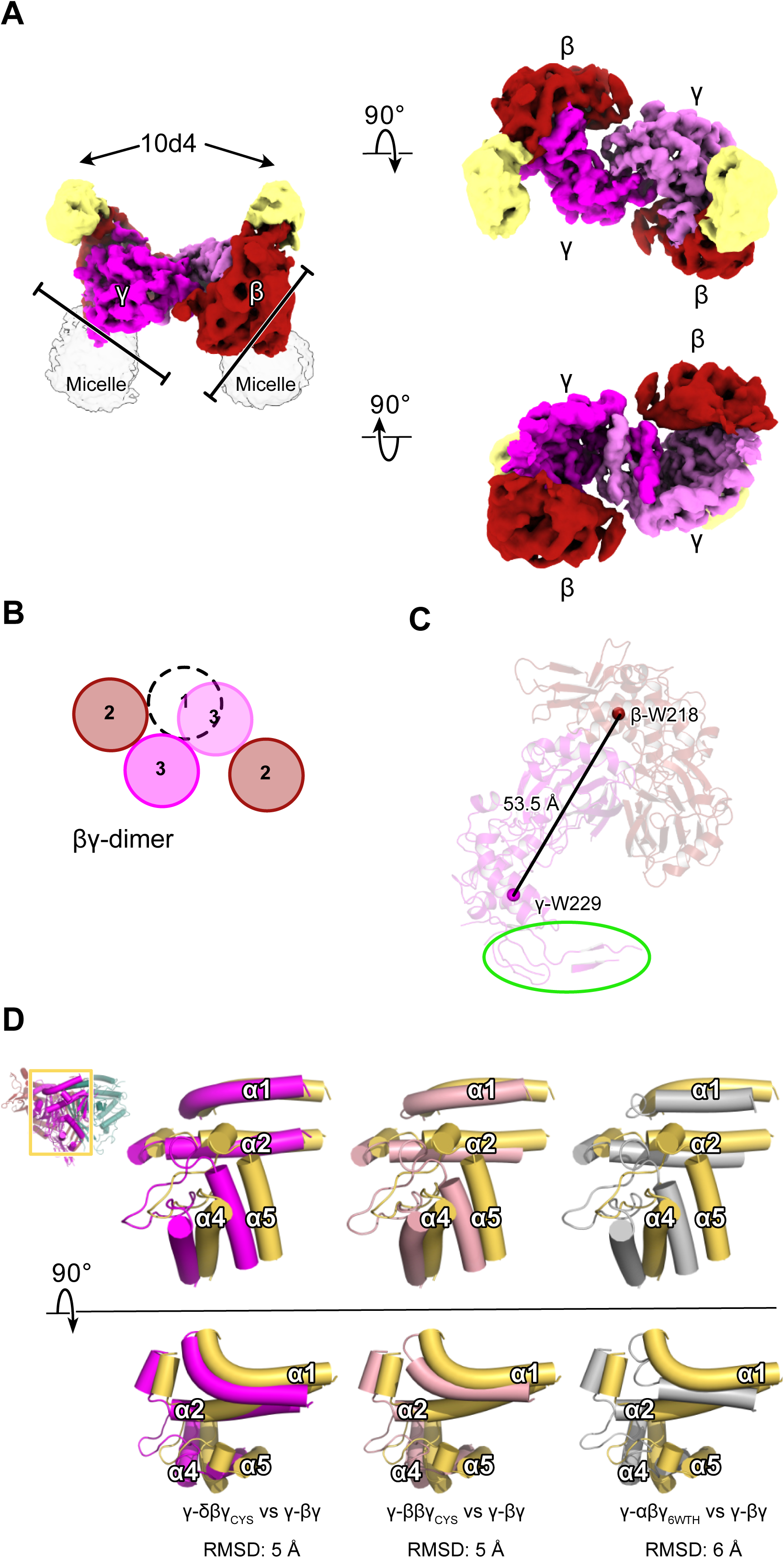
Heteromeric ENaC assembly intermediate shows β and γ assemble as dimers. A. Cryo-EM map of the βγ-dimer with subunits and Fab colored as in figure 2. The second γ subunit is colored in light pink for clarity. Solid bars indicate the direction of the putative membrane plane. B. Schematic illustration of the dimer-dimer interaction shown in A with the missing position 1 subunit shown as dashed circle. C. Cartoon representation of the βγ dimer with distances of the Cα atoms of conserved tryptophans in the α2 helices. The highlighted region in green is the γ-GRIP portion that is not resolved in ENaC trimers. This region mediates extensive interdimer contacts. D. Inset: An overall view of the δβγCYS extracellular domain in cartoon representation. The region highlighted with a yellow rectangle includes the finger and thumb domains of γ. Close-up views of the boxed region showing comparison of the α1, α2, α4, and α5 positions in the finger and thumb domains of γ from trimers to the γ from βγ-dimer. The γ from trimers are colored as in figures 4 and 5, while γ from the dimer is colored yellow. All helices are shown in cylinders. See also Figure S7.

The βγ dimer provides an opportunity to investigate assembly in ENaC. Although there appears to be a loop emanating from the γ GRIP domain that could obstruct access to position 1, we speculate that this loop is present in our data primarily because it is stabilized by its interaction with the second βγ dimer. Assuming this loop is flexible and can move aside when a position 1 subunit joins to form the trimer, we focused on one dimer for the rest of the structural analysis. Implementing symmetry expansion to enhance map quality of one dimer, βγ, we achieved a reported resolution of 3.7 Å (**Figure S7**). The map quality improvement enabled us to trace the main chain in regions that were previously unmodeled in the γ-GRIP domain (**Figure 7C, S7**). Using this model, we measured the distance between the β and γ Cα TriTrp, and this distance was approximately 53.5 Å, which is longer than the distances measured in the αβγ_6WTH_ and ββγ_CYS_ trimers but slightly shorter than the δβγ_CYS_ (**Figure 3A-C**). This suggests that to accommodate a subunit in position 1, the γ finger must rearrange (**Figure 3A**). We tested this hypothesis by aligning the position 2 β subunits of all trimers and the βγ dimer and compared the positions of the finger and thumb domains of γ. We found that the finger and thumb domains of γ subunits from all three trimers adopt different positions when compared to the γ-βγ. With a calculated RMSD of 6 Å, the finger and thumb of γ-αβγ_6WTH_ are most different from γ-βγ (**Figure 7D**).

Examining the three trimers has allowed us to measure the conformational changes the γ subunit undergoes to accommodate various subunits in position 1. The predominant conformational shifts occurring in the finger domain suggest that, while not as rigid as β, γ in position 3 accommodates α, β, and δ subunits. It is worth noting that the pore and cytosolic domains may also influence subunit stoichiometry and trimer conformation. However, the lack of information in these areas limits our understanding of their impact on trimeric assembly. Nevertheless, the observed trimers and dimer complexes in our study offer valuable structural insights into the diverse assemblies of ENaC.

## Discussion

The trimeric organization of ENaC is evident in three ENaC trimer structures: the previously determined structure of human αβγ trimer, and in this study, the δβγ_CYS_, and ββγ_CYS_ trimers. This display of heteromeric assembly suggests that these ion channels can tune Na^+^ reabsorption by leveraging the contributions of individual subunits. The selective isolation method implemented in this research prioritizes γ-containing assemblies, potentially overlooking other ENaC proteins with different subunit compositions.^58,66–69^ Interestingly, this bias towards γ-containing proteins did not result in observable capture of homomeric assemblies or proteins with two γ subunits. This might indicate limitations in the recombinant system used or unique properties of the γ subunit that inhibit assembly with multiple γ subunits.

This study describes a list of observations that either ground decades of functional experiments in three dimension or unveil novel insights that will serve as foundations for future experiments to understand ENaC function. First, the structure of the human δβγ offers insights into the δ-containing heteromeric assembly, elucidating the molecular basis for subunit-dependent functional differences between δβγ and αβγ ENaC. A consistent pattern emerges in these diverse trimers, where β and γ subunits consistently occupy positions 2 and 3, while δ or α takes position 1. The impact of the subunit in position 1 is evident through variations in TriTrp distances; the presence of δ leads to an expanded TriTrp triangle compared to the trimer containing α. Second, while the functional relevance of the ββγ trimer is currently obscured by its failure to traffic to the plasma membrane in HEK cells, the obtained structure provides the first evidence of how a trimer can form containing only β and γ subunits. Understanding the functional impact of the β subunit in position 1 proves challenging. However, we can speculate a possible functional consequence based on the conformations of the GRIP domains leading us to the third key observation: the protease-sensitive GRIP conformation. An examination of the γ-GRIP domain suggests that the conformation in αβγ, sensitive to proteases, differs from that in δβγ, which shows protease resistance according to previous studies.^26,28^ If we were to infer the functional consequences based on the observed conformations in the γ-GRIP domain, ββγ is likely to exhibit a protease response akin to δβγ. This proposed lack of protease sensitivity is consistent with recent findings examining the response of ENaC channels with diverse subunit compositions. In this study, ENaC responds to shear stress, even in cases where only β and γ subunits are present.^58^

Fourth, the homogeneous conformation observed in the β subunits in position 2 of all three trimers is striking and suggests a scaffolding role of β when occupying position 2. It is likely that the role of position 1 subunit is to define the mode of stimulus to which the channels respond. Moreover, the presence of γ in position 3 ultimately regulates the degree of the response. This aligns with findings from multiple studies on αβγ and its response to proteases, where full cleavage of the γ subunit substantially increases *P_o_* to 1.^55,70–72^ Fifth, this study marks the initial characterization of human δβγ modulation by Zn^2+^. The opposite responses of δβγ and αβγ to specific concentrations of Zn^2+^ underscore the functional diversity of ENaC. Obtaining a structure of δβγ and αβγ in the presence of Zn^2+^ could shed light on how divalent cations modulate ENaC activity.

Sixth, the presence of a molecule in the palm domains of ββγ_CYS_ and δβγ_CYS_ is surprising, offering a new opportunity to explore the function of ENaC. In δβγ_CYS_, the molecule feature is not as well-defined as in ββγ_CYS_, correlating with the weak map feature of the δ palm domain. A possible explanation could be that there is a mixed population of δβγ_CYS_ complexes, bound or unbound to the molecule, resulting in conformational heterogeneity and poor map quality in this region. The molecule seems to favor binding with the ββγ_CYS_resulting in an overall well-resolved 3D map of the complex. The lower palm region in αβγ is also well-defined and does not display a similar feature suggesting that this peptide-like molecule does not favorably bind to the αβγ trimer.^36,37^ Perhaps, the compact nature of αβγ as demonstrated in its TriTrp distances may hinder access to this unidentified molecule. The positions of the β strands in the lower palm domain in αβγ is incompatible with peptide binding. For the molecule to bind, rearrangement of the β sheets would be required.

Lastly, the βγ dimer implies the potential assembly of the two subunits. Previous studies indicate abundant expression of β and γ subunits in the kidneys at the protein level.^60,62,65^ Whether these subunits exist in monomeric or dimeric forms remains unclear. However, our findings suggest that they have the capacity to assemble into dimers, possibly serving as assembly intermediates. Similar intermediates have been previously observed in pentameric cys-loop receptors.^73^ In cys-loop receptors, a subset of particles derived from a mouse tissue sample consisting of only three and four subunits was identified, suggesting the presence of assembly intermediates.^73^ Therefore, in the context of ENaC, it is plausible to consider assembly intermediates in more dynamic processes akin to processes observed in the kidney. In kidneys, this assembly intermediate could efficiently respond to fluctuations in salt intake by preemptively having dimers ready. Completing the trimer by inserting the third subunit in position 1 creates a functional trimer ready for Na^+^ reabsorption.^43,65^ We acknowledge that the lack of information on the pore domain prevents assigning a functional state to our structures. Nevertheless, the three structures outlined in this study lay the groundwork for future investigations, enabling us to further explore the molecular mechanisms governing the diverse responses driven by the compositional differences in ENaC channels.

## Acknowledgements

We thank members of the Baconguis and Gouaux labs for discussion on this study. We especially appreciate the feedback from Arpita Bharadwaj and Kimberly Hartfield on the manuscript. A portion of this research was supported by NIH grant U24GM129547 and performed at the Pacific Northwest Center for Cryo-EM at Oregon Health & Sciences University (OHSU) and accessed through EMSL (grid.436923.9), a DOE Office of Science User Facility sponsored by the Office of Biological and Environmental Research. Initial experiments included in this study were supported by a grant from the NIH to I.B. (GM138862). Follow-up experiments were supported by a grant from the Cystic Fibrosis Foundation to I.B. (BACONG22G0). A.H. was supported by the National Science Foundation Graduate Research Fellowship Program (GVPRS0015B4).

## Author contributions

A.H and I.B. designed the project. A.H. carried out expression, purification, sample preparation for cryo-EM analysis, cryo-EM processing, model building, imaging, and electrophysiology experiments. A.H. and I.B. wrote the manuscript.

## Supplemental information titles and legends

Document S1. Figures S1-S7 and Tables S1-S2.

## STAR Methods

### Key resources table

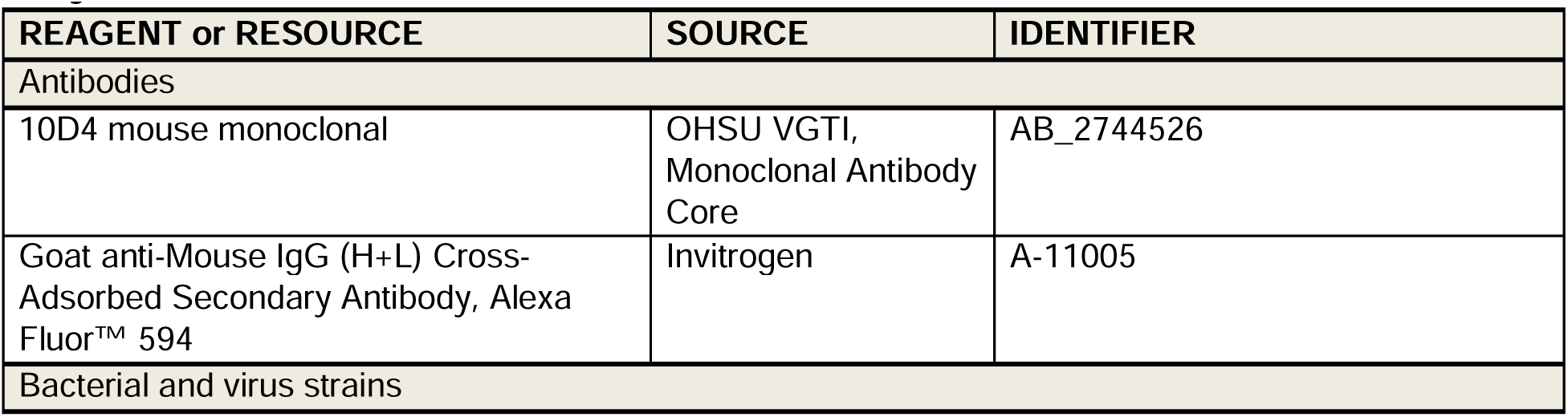
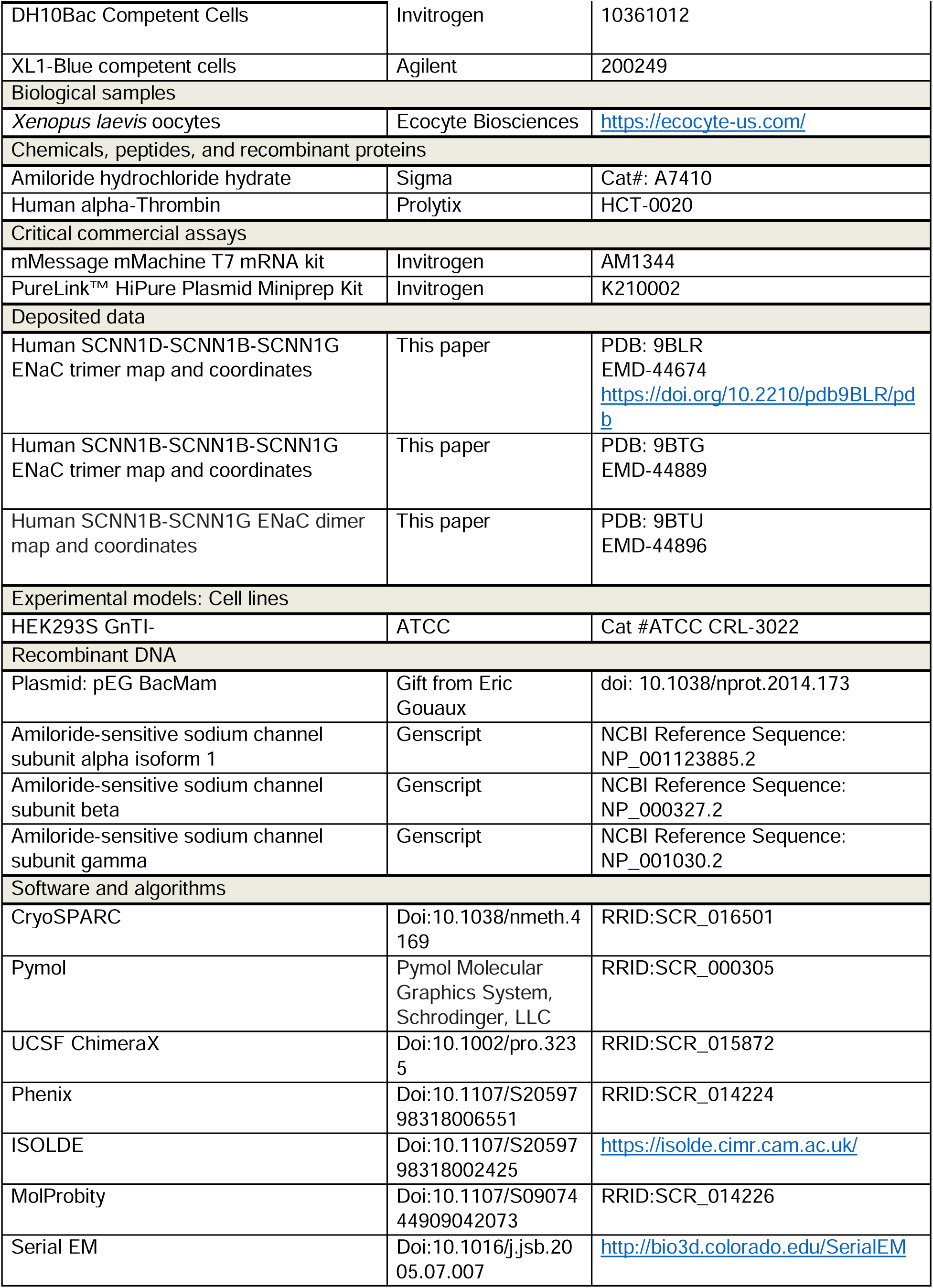
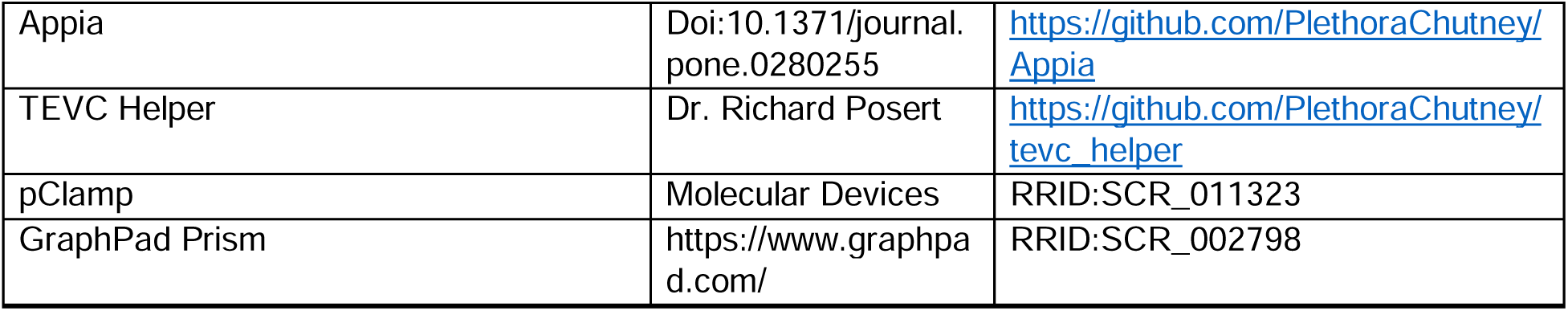

### Resource availability

#### Lead contact

Further information and requests for resources and reagents should be directed to and will be fulfilled by the lead contact, Isabelle Baconguis.

#### Materials availability

For any correspondence or requests related to the materials used in this study, please contact the lead contact.

#### Data and code availability

- Cryo-EM maps and atomic models for δβγ_CYS_, ββγ_CYS_, and βγ_Dimer_ have been deposited in the Protein Data Bank and Electron Microscopy Data Bank and are publicly available as of the date of publication. The PDB IDs are 9BLR, 9BTG, and 9BTU. The EMDB IDs are EMD-44674, EMD-44889, and EMD-44896.
- This study does not employ any original code.
- Any additional information required to reanalyze the data reported in this work is available from the lead contact upon request.

#### Experimental model and study participant details

The cells used for generating baculovirus are Sf9 cells (*Spodoptera frugiperda,* Cat# CRL-1711) and cultured in 27° C. HEK293S GnTI-suspension cells used for expression of all ENaC proteins in this study were obtained from the ATCC (Cat# CRL-3022) and cultured at 37°C and 8% CO_2_.

### Method details

#### Construct design

The gene encoding the wild-type human δ ENaC subunit was codon optimized, synthesized, and cloned in the pEG BacMam vector by Genscript.^74^ The wild-type sequences of the human β and γ subunits from previous structural studies of human αβγ were used. The γ construct consists of an octa-histidine tag, eGFP, and a Thrombin cleavage site at the N-terminus.^37^ ENaC subunit mutants were also generated by Genscript by site-directed mutagenesis to introduce mutations in cysteine residues within the preTM1 region of each ENaC subunit, along with the R138A mutation in the γ subunit to render it furin-resistant. The δ subunit, along with its own CMV promoter, was then isolated as a cassette using restriction enzymes PmeI and AvrII and integrated into a pEG BacMam vector containing the N-terminally eGFP tagged γ subunit. This process was repeated with the β subunit so all three genes encoding the δ, β, and γ subunits of ENaC were present in a single plasmid, each with its own CMV promoter. The presence of all ENaC subunit genes were verified by agarose gel electrophoresis following selected restriction enzyme digestion and DNA sequencing. Subsequently, these plasmids were employed to generate bacmid, followed by the production of P1 and P2 viruses for the expression of ENaC.^74^ The generated viruses were used for various purposes, including electrophysiology studies, expression analysis, and purification.

#### Small-scale expression and solubilization

For expression optimization, we performed small-scale expression by infecting 20 mL suspension cultures of HEK293 GnTI-at a density of 3x10^6^ cells/mL with baculovirus at an MOI of 2. Baculovirus generation is described in the Construct design section. Following an incubation at 37°C for 8 hours, 100μM amiloride and 10μM sodium butyrate final concentration were added, and the cells were transferred to an incubator set at 30°C for protein expression over 24 hours. Then 2 mL aliquots of cells were centrifuged at 5000*g* for 10 minutes. The resulting pellets were flash-frozen and stored at 80°C. On the day of the experiment, the frozen samples were thawed on ice, and the pellets were washed with 1 mL of HEPES Buffered Saline (pH7.4). Then, 150μl of either DDM solubilization buffer (20 mM n-dodecyl-β-D-maltoside, 3 mM cholesteryl hemisuccinate, 200 mM NaCl, 20 mM HEPES, pH 7.4 and Thermo Fisher Halt Protease Inhibitor Cocktail at a volume ratio of 1:100) or digitonin solubilization buffer (7% digitonin, 200 mM NaCl, 20 mM HEPES pH 7.4 and Thermo Fisher Halt Protease Inhibitor Cocktail at a volume ratio of 1:100) was used to resuspend the washed pellet, which were then allowed to nutate at 4°C for 1 hour. Samples were then spun at 71,680*g* for 20 minutes. The resulting supernatant was loaded into 0.22μm filter tubes and spun at 4000*g* for 5 minutes at 4°C. Afterward, the samples underwent another round of centrifugation for 40 minutes at 71,680*g* before proceeding with FSEC or western blot analysis.

#### Patch-Clamp Electrophysiology

Suspension-adapted HEK293 GnTI-cells were prepared at a density of 3x10^6^ cells/mL and infected with baculovirus, generated as described in Construct design section, at a multiplicity of infection (MOI) of 2. Cells were incubated at 37°C with agitation for 12 hours. Subsequently, amiloride and sodium butyrate were added with a final concentration of 100 μM and 10μM, respectively. The cells were transferred to an incubator set at 30°C. Twenty-four hours post-infection, the cells were plated on glass cover slips in DMEM with 10% FBS at a volume ratio of 1:4 for cells to media.

Whole-cell patch-clamp recordings were carried out 12 hours post-plating the cells in DMEM. Patch pipettes were pulled and polished to 2–3 MΩ resistance using the Sutter Instrument P-97 and Narishige MF-830 Microforge. Pipettes were filled with an internal solution containing 150 mM KCl, 2 mM MgCl_2_, 5 mM EGTA, and 10 mM HEPES (pH 7.4). For the recording of whole-cell amiloride-sensitive currents, the cells were placed in a bath solution containing 100 μM amiloride, 150 mM NaCl, 2 mM MgCl_2_, 2 mM CaCl_2_, and buffered with 10 mM HEPES (pH 7.4). To measure amiloride-sensitive currents, cells were perfused with an external solution that was the same as the bath solution followed by a second external solution lacking amiloride. The amplitude of the amiloride-sensitive current was determined by the difference between conditions with and without amiloride. For bi-ionic experiments, NaCl was replaced with 150 mM LiCl or 150 mM KCl in the external solution. To generate current-voltage (I-V) trace, currents were recorded while the cell was held at voltage potentials ranging from -80mV to 80mV in 20mV increments while the external solutions containing Na^+^, Li^+^, or K^+^ with or without amiloride were perfused onto the cell. Data was acquired with the Axon Instruments Molecular Devices Axoclamp 200B and Axon DigiData 1550B. Data was analyzed with pClamp, GraphPad Prism, and ABF Plotter.^75^

#### Two-electrode voltage-clamp electrophysiology

Individual wild-type ENaC subunit constructs were cloned into the pcDNA3.1 vector modified to contain a T7 promoter by GenScript. These vectors were linearized and transcribed to generate mRNA using Invitrogen’s mMessage mMachine T7 mRNA kit. Oocytes from Ecocyte Bio Science were injected with a total volume of 50 nL containing 1.5 ng of each ENaC subunit. Following injection, oocytes were then incubated at 19°C in MBSH (88 mM NaCl, 1 mM KCl, 2.4 mM NaHCO_3_, 10 mM HEPES (pH 7.4), 0.33 mM Ca(NO_3_)_2_, 0.41 mM CaCl_2_, 0.82 mM MgSO_4_) and supplemented with 150 µg/mL Gentamycin, 250 µg/mL Amikacin, and 100 µM Amiloride. ENaC activity was recorded 18-24 hours post-injection using electrodes filled with 3 M KCl. Bath solutions used were variations of Ringer solution containing 110mM of cation, 1mM KCl, 1.8 mM CaCl_2_, 5 mM HEPES (pH 7.4). Data was acquired with the Axon CNS Molecular Devices Axoclamp 900A and Digidata 1440A. Data was analyzed with pClamp, GraphPad Prism, and TEVC Helper.^75^

#### Large-scale ENaC expression and purification

Suspension-adapted HEK293 GnTI-cells were prepared at a density of 3x10^6^ cells/mL and infected with baculovirus. Following an incubation at 37°C for 8 hours, 100 μM amiloride and 10 μM sodium butyrate were added, and the cells were transferred to an incubator set at 30°C for protein expression over 72 hours. Post-infection, the cells were spun at 5000*g* for 20 minutes, the supernatant was discarded, and the pellet was flash frozen and stored at -80°C. To begin the purification process, the frozen cell pellet was slowly thawed on ice followed by homogenization in a solubilization buffer consisting of 1% digitonin, 20 mM HEPES pH 7.2, 200 mM NaCl, 2 tablets of Thermo Fisher Protease Inhibitor Cocktail, 2 mM ATP, and 2 mM MgSO_4_. The solubilization was achieved by stirring the solution at 350 rpm at 4°C for 1.5 hours, followed by centrifugation at 125,440*g* for 1 hour to collect the supernatant.

The supernatant was then passed through a GFP Nanobody (GNB) resin packed into an XK 16/40 column for binding. The column underwent successive washing steps: first with 5 column volumes of buffer A containing 0.1% digitonin, 200 mM NaCl, and 20 mM HEPES pH 7.4, then with 2 column volumes of Buffer A supplemented with 2 mM ATP and 2 mM MgCl_2_, followed by an additional 5 column volumes of Buffer A. For elution, the column was incubated in Buffer A supplemented with 5 mM CaCl_2_ and 34 μg/mL of Thrombin for 15 minutes at room temperature, from which one column volume separated into fractions were collected. After an additional 30-minute incubation with Thrombin, an additional column volume was collected in fractions. Fractions were run on SDS-PAGE and A260/A280 was measured on a Thermo Scientific NanoDrop One. Samples containing high levels of protein were concentrated with Millipore Amicon Ultra-15 Centrifugal Filters, 100 kDa molecular weight cut-off. Subsequently, the 10d4 Fab was added to the concentrated sample at a molar ratio of 1.5:1 of Fab:ENaC before undergoing centrifugation at 71,680*g* for 40 minutes. This step was followed by size-exclusion chromatography using a Superose 6 increase 10/300 GL column. Peak fractions were collected. Samples obtained at various purification steps were subjected to examination by SDS-PAGE and assayed by FSEC^40^. SEC fractions verified to contain ENaC by SDS-PAGE and FSEC were pooled and concentrated to 3-4 mg/mL for grid preparation.

#### Cryo-EM data aquisition and analysis

The concentrated sample was subjected to centrifugation at 71,680*g* for 10 minutes at 4°C followed by addition of fluorinated octyl maltoside to 10 µM. Then, 3 µl of the sample was applied to glow-discharged (15 mA for 60 seconds) Quantifoil holey carbon grids (Au 2 µm/1 µm 200 mesh). The grid was manually blotted followed by a second application of 3 µl, which was then blotted using a Vitrobot Mark IV (FEI). The Vitrobot blot parameters were set to a wait time of 0s, blot time of 2s, and a blot force of 1 with the temperature maintained at 22°C and humidity at 70%. Grids were then flash frozen in liquid ethane.

Data was collected on a 300_KeV Titan Krios equipped with K3 direct electron detector at the Pacific Northwest Cryo-EM Center (PNCC). Acquisition was automated using SerialEM to find holes with suitable ice thickness using the hole finder feature and combined to produce multishot–multihole targets.^76,77^ A total dose of 50_e^−^/Å^2^ was divided into 50 frames, with a pixel size of 0.8015 Å/pix (0.40075 Å/pix with super resolution) and a defocus range of -0.8 to -2.5 µm. During data collection, CryoSPARC Live was employed initially to handle preliminary processing and provide an overview of the data collection progress.^78^ Once the data collection finished, the acquired movies were imported into CryoSPARC for further processing, including motion correction and CTF estimation. Particles were initially picked using blob picker and cleaned using rounds of 2D classification. The selected particles were used for *ab-initio* reconstruction to generate templates for subsequent template-based particle picking. Template-picked particles underwent additional rounds of refinement including heterogeneous refinement and 3D classification aimed at producing a set of “good” classes representing recognizable ENaC classes. Subsequently, the final particle set was then subjected to local refinement using masks to exclude Fab regions. Multiple rounds of local refinement were performed using different masks to optimize the final reconstructions.^78–80^ Finally, the generated maps underwent manual inspection to ensure their quality met the standards necessary for model-building. Additionally, this inspection aimed to assess any limitations that may affect interpretation in subsequent structure analysis.

#### Model building

The preliminary models of the β and γ subunits derived from PDB 6WTH were initially fitted into the experimental maps. Conversely, for the δ subunit, its initial model was generated utilizing AlphaFold2.^57^ Before docking into the map, both the pore and cytosolic domains were removed. Following this, all three subunit models underwent rigid body fitting, accompanied by the removal of loops within disordered regions using Coot.^81^ A series of iterative model-building employing Isolde and Coot, along with real-space refinement in Phenix, were conducted until the final models were considered to have satisfactory stereochemistry, as defined by MolProbity.^81–85^ The final model of the δβγ_CYS_ trimer contains residues 126-516 with regions unmodeled in the α2-β4 loop, α6-β11 loop, and the GRIP domain for the δ subunit, residues 78-512 with a region unmodeled in the GRIP domain for the β subunit, and residues 80-521 with regions unmodeled in GRIP domain for the γ subunit. The ββγ_CYS_ trimer model contains residues 78-512 with regions unmodeled in the GRIP and knuckle for the β^1^ and β^2^ subunits, and residues 79-522 with regions unmodeled in the GRIP domain of the γ subunit. The βγ contains residues 80-510 with a region unmodeled in the GRIP domain β subunit, and residues 85-516 with regions unmodeled in the GRIP domain for the γ subunit. For structure analysis, angle_between_helices and center_of_mass plugin function in PyMOL Molecular Graphics System (Version 3.0 Schrödinger, LLC) and the rmsd command in ChimeraX were implemented.^86–88^ To generate figures for this study, both PyMOL and ChimeraX were used.

#### Immunofluorescence Staining

HEK293 GnTI-cells, infected at 3x10^6^ cells/ml, were plated in DMEM supplemented with 10% FBS on a glass bottom 35mm dish resulting in a final cell density of 0.45 x10^6^ cells/ml. The cells were allowed to settle at 30°C for 5 hours. Subsequently, they were rinsed with 1mL PBS and fixed with 4% PFA in PBS for 10 minutes at room temperature with gentle agitation. After fixation, the cells were rinsed twice with 1mL PBS. For permeabilization, 0.1% Triton X-100 in PBS was added and incubated for 20 minutes at room temperature on a nutator. Following this step, the cells were washed twice with PBS. To reduce non-specific binding, 3% BSA in PBS was applied to all samples and left at 4°C overnight. The next morning, the monoclonal antibody 10d4, directed against the β subunit, was applied at a concentration of 4µg/ml in 3% BSA PBS and incubated for 2 hours at room temperature while nutating. Subsequently, the cells were washed with PBS three times for 5 minutes each. For fluorescent labeling, Alexa Fluor 594 goat anti-mouse antibody, diluted in 3% BSA in PBS at a volume ratio of 1:1000 of antibody to total volume, was added for 1 hour at room temperature on a nutator while being covered in foil to prevent photobleaching. After staining, the cells were washed three times with PBS for 15 minutes each while being kept covered in foil. Images were collected with a Leica DMi8 inverted microscope and analyzed with Leica Application Suite Version 4.12.0.

## Supplementary figure legends

**Figure S1. Design and optimization of δβγ constructs for structural investigations by cryo-EM, related to Figure 1**.

**A.** Schematic illustration of the three δβγ constructs: δβγ_WT_, δβγ_R138A_, and δβγ_CYS_. **B.** Current-voltage experiment demonstrating that δβγ_WT_ is Na^+^-selective, permeable to Li^+^, and impermeable to K^+^ when expressed in *Xenopus laevis* oocytes. Voltage potential range used for the experiment is -80 mV to 80 mV in 20 mV increments. The external solutions contained equimolar concentration of Na^+^, Li^+^, and K^+^. The pipettes contained 3M KCl. Each point is represented as mean ± SEM (n = 5). **C.** Dose-response of δβγ to Zn^2+^. Current ratios were determined by measuring current amplitudes before and after application of Zn^2+^. Data are represented as mean ± SEM (n=12). **D.** Dose-response of αβγ to Zn^2+^ using the same concentration range as in C. Data are represented as mean ± SEM (n=3). **E.** Normalized FSEC traces comparing the biochemical behavior of purified δβγ_WT_ and αβγ_WT_. Traces were normalized to the height of the peak at 14 mL.

**Figure S2. Three-dimensional reconstruction details of δβγ_CYS_ and ββγ_CYS_, related to Figure 2**.

**A.** Overview of the cryo-EM data processing workflow that revealed two trimeric complexes: δβγ_CYS_ and ββγ_CYS._ **B.** Cryo-EM maps of the subunits in δβγ_CYS_. The glycosylation sites are colored yellow, and the subunits are colored as in Figure 2. **C-H.** Fourier Shell Correlation curves, local resolution maps, and angular distribution plots for δβγ_CYS_ and ββγ_CYS_.

**Figure S3. Cryo-EM maps of the gating domains in δβγ_CYS_, related to Figure 3**.

**A-C.** Cryo-EM maps of the finger, thumb, and knuckle domains of δ (A), β (B), and γ (C) subunits and the corresponding models.

**Figure S4. Comparison of the finger, thumb, and knuckle domains and interfaces of subunits, related to Figure 4**.

**A.** Superposition of the β subunits occupying position 2 from δβγ_CYS_, ββγ_CYS_, and αβγ_6WTH_. **B.** The close-up views of the boxed area in positions 2 and 3 from figure 4E are displayed. Sidechains of residues at corresponding positions in β and γ from ββγ_CYS_ and αβγ_6WTH_ are shown in stick form to illustrate the orientation of their side chains relative to the pore axis, symbolized as a black circle in the upper left corner of the view. The black circle marks the relative direction of the pore axis position. **C.** A view of the finger domain showing residues that form contacts between β6-β7 loop and the helical segments. Sidechains are shown in sticks and dots representation to provide visualization of sidechain packing. **D.** Superposition of the position 1 subunits and comparison of the thumb domains. **E.** Views of the position 1/position 2 and position 2/position 3 interfaces in ββγ_CYS_ and αβγ_6WTH_.

**Figure S5. The γ subunit finger domain rearranges in the presence of δ and β in position 1, related to Figure 5**.

**A.** Cryo-EM map of the β6-β7 loop in ββγ_CYS_ and the corresponding residues. **B.** Superposition of the γ subunits from δβγ_CYS_, ββγ_CYS_, and αβγ_6WTH_. **C.** Cryo-EM maps of δβγ_CYS_, ββγ_CYS_, and αβγ_6WTH_. The γ-GRIP domains are colored blue while the rest of the map is shown in gray. The new map features observed in δβγ_CYS_ and ββγ_CYS_ are colored orange. **D.** The model generated by AlphaFold2 for δβγ reveals that the γ-GRIP region consists of several β strands. Among these strands, one positioned close to the upper part of the GRIP segment is likely the novel feature detected in the δβγ_CYS_ and ββγ_CYS_ maps.

**Figure S6. Cryo-EM analysis, functional, and biochemical characterization of ββγCYS, related to Figure 6**.

**A.** Model of ββγ_CYS_. The region forming contacts with the peptide-like feature in the palm domain is highlighted. **B.** Close-up view of the colored region in A. **C, D.** Current-voltage relationship of cells infected with β and γ baculovirus (C) and uninfected cells (D). **E, F.** Immunofluorescence staining of unpermeabilized (E) and permeabilized (F) cells infected with δβγ (left), βγ (middle), or left uninfected (right). Surface expression of ENaC was monitored using the 10d4 monoclonal antibody directed against the β subunit. A secondary antibody labeled with Alexa Fluor 594 was used for fluorescence labeling.

**Figure S7. Cryo-EM analysis of the pop1 peak from δβγ_WT_-expressing HEK cells, related to Figure 7**.

**A.** Illustration of the cryo-EM workflow used to generate the final βγ dimer map. **B-D.** FSC curves, local resolution estimation, and angular distribution plot of βγ dimer.

## Supplementary figure legends

**Table S1. Cryo-EM data collection, refinement, and validation statistics**

**Table S2. Root-mean-square deviation of C**α **atoms in each subunit domains**

Using the rmsd command in ChimeraX, RMSD values were calculated by comparing the Cα positions of each subunit domain between two subunits. The three columns represent RMSD values obtained through three different superpositioning methods: individual subunit superposition, trimer superposition, and trimer superposition using the position 2 β subunit as a reference. The values are color-coded according to the subunit domain: the upper palm, knuckle, β-ball, finger, and thumb domains are colored yellow, cyan, orange, purple, and green, respectively.

