## Supplementary figures and images for "Structural Insights into Subunit-Dependent Functional Regulation in Epithelial Sodium Channels"

### Figure S1

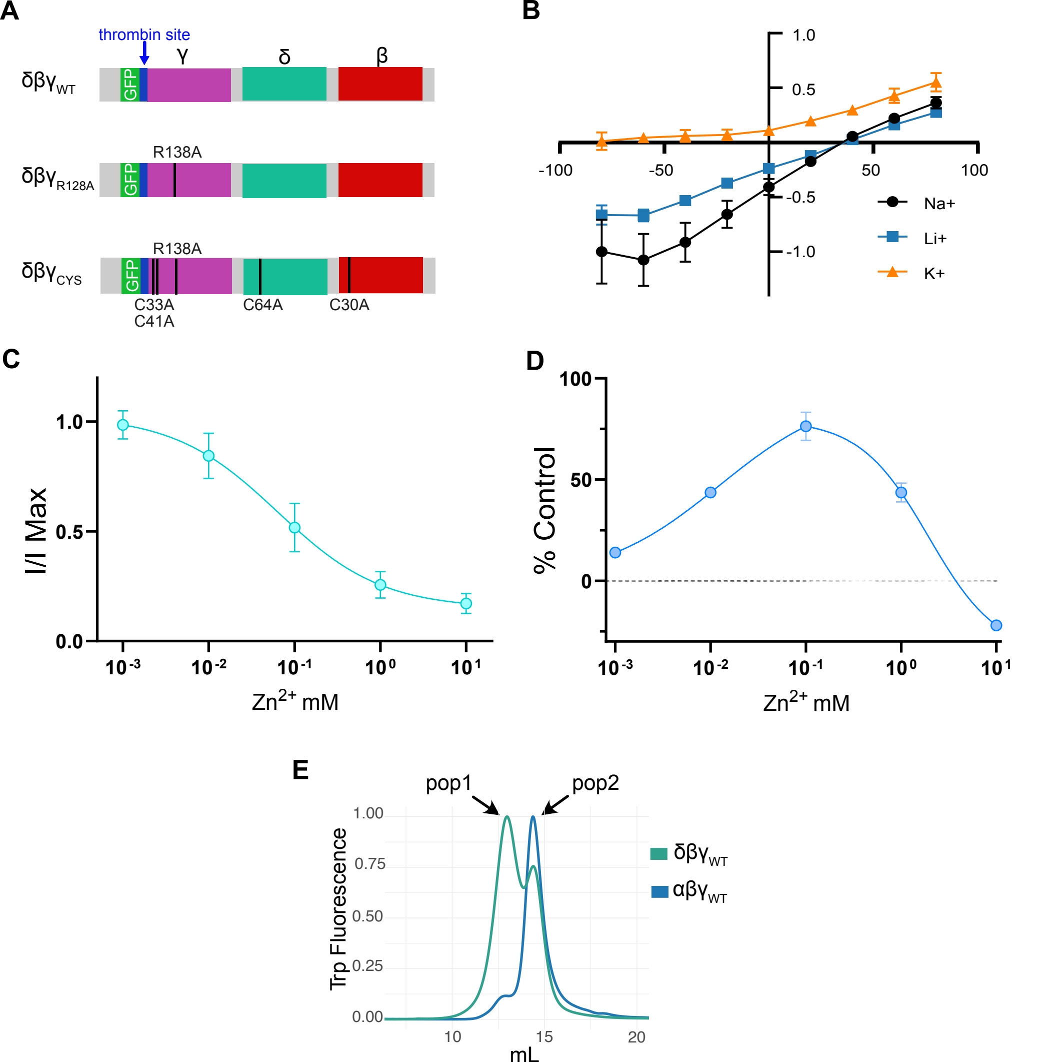

### Figure S2

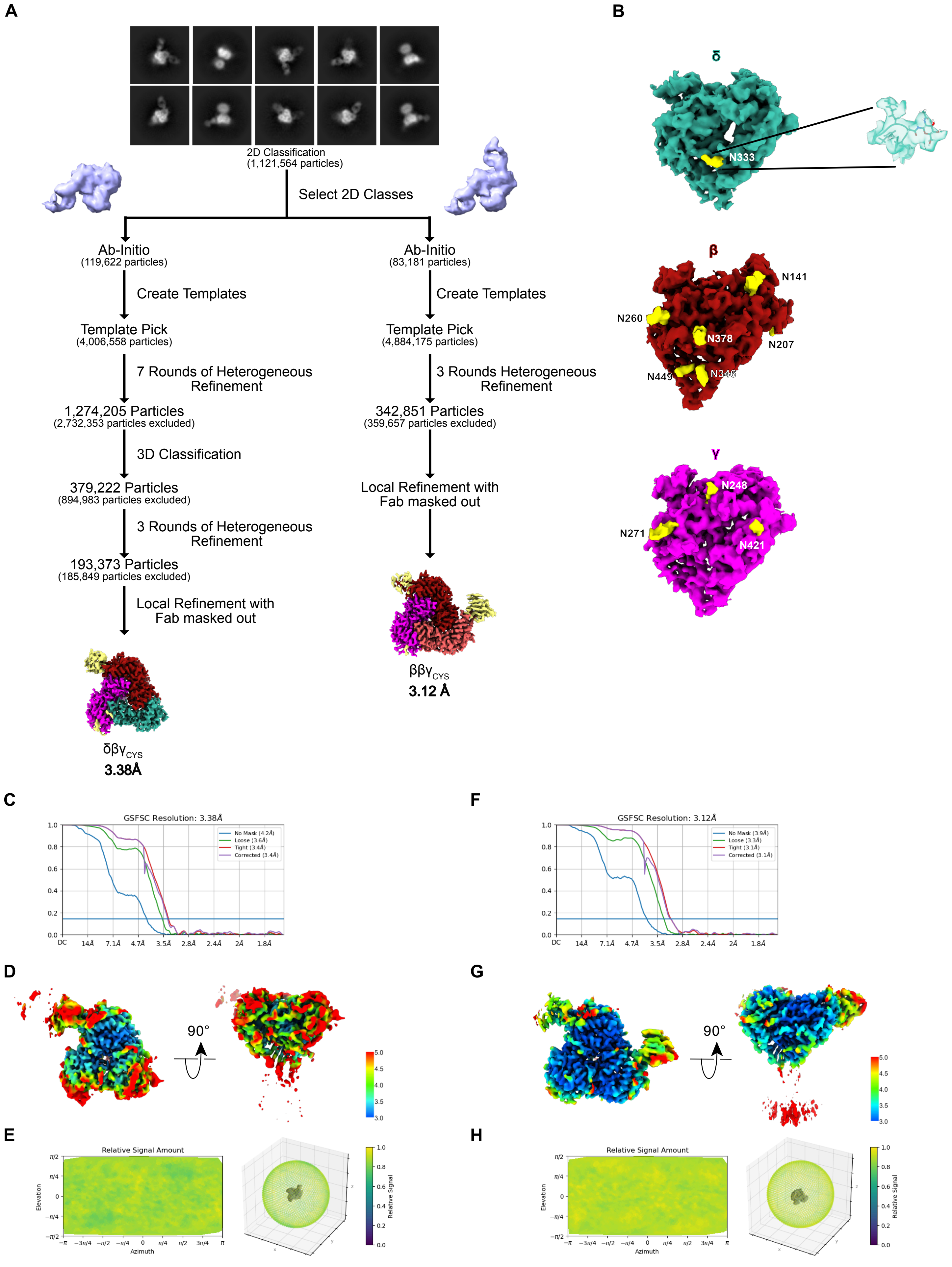

### Figure S3

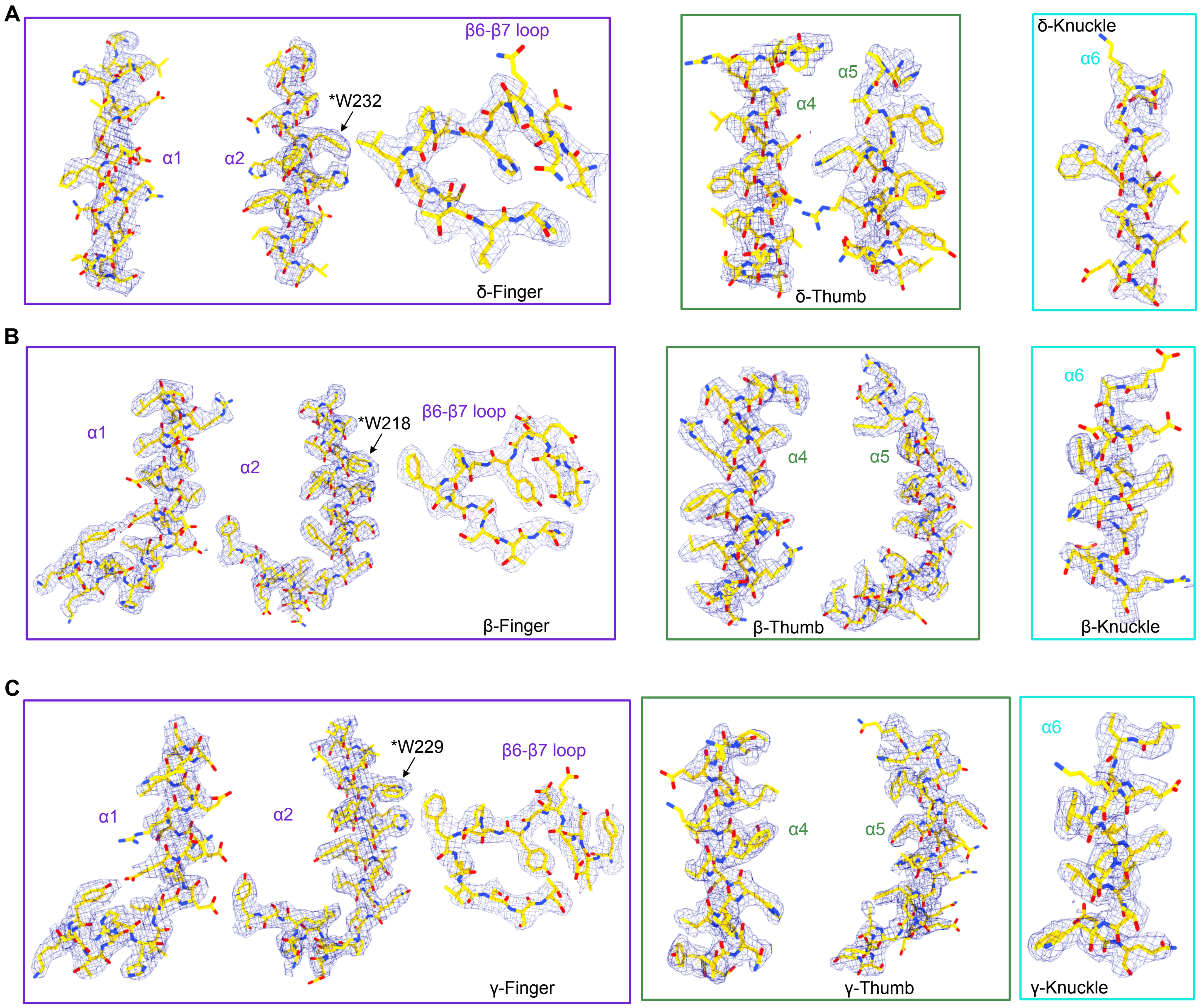

### Figure S4

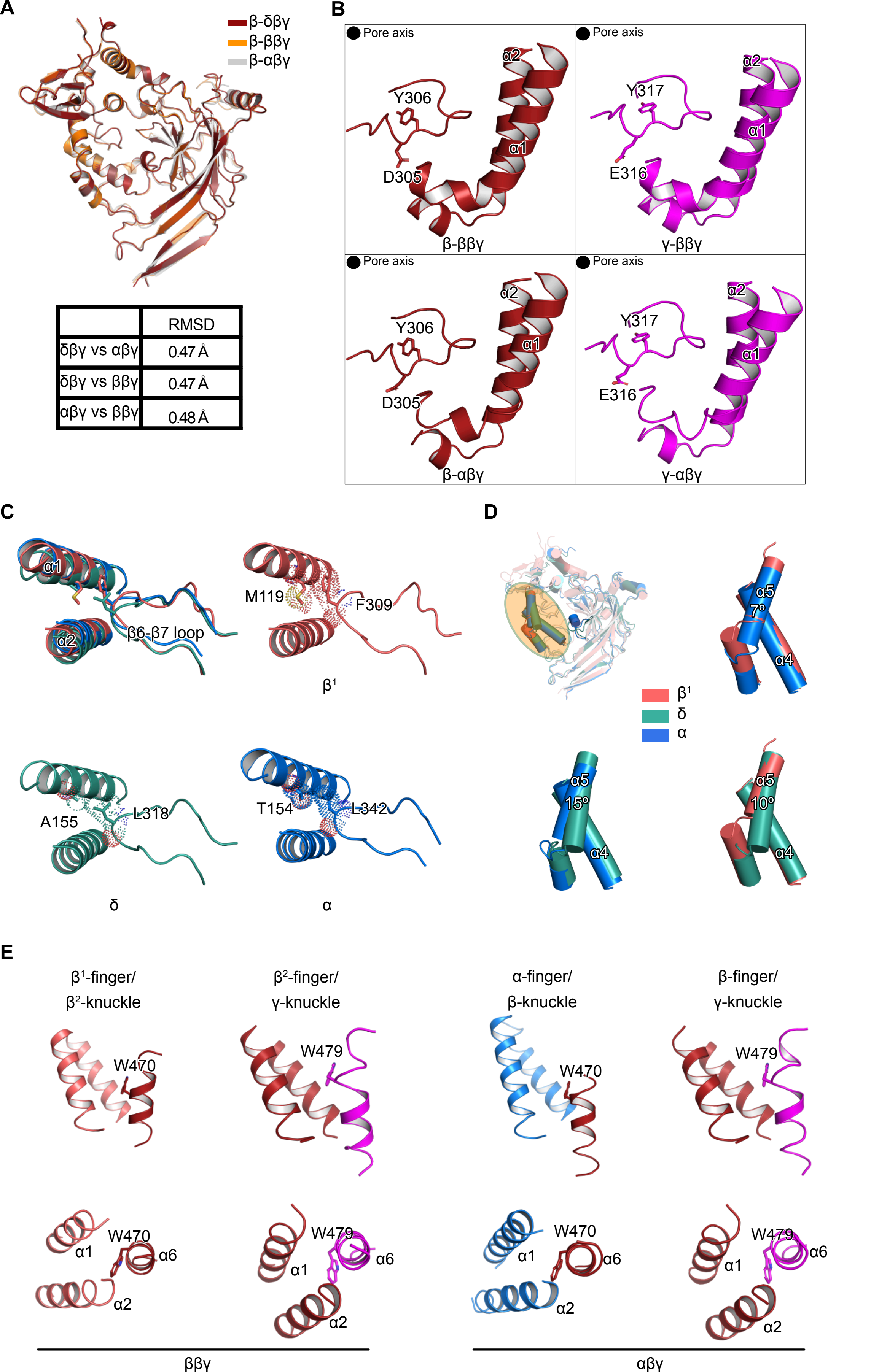


| | RMSD |
|---|---|
| δβγ vs αβγ | 0.47 Å |
| δβγ vs ββγ | 0.47 Å |
| αβγ vs ββγ | 0.48 Å |

### Figure S5

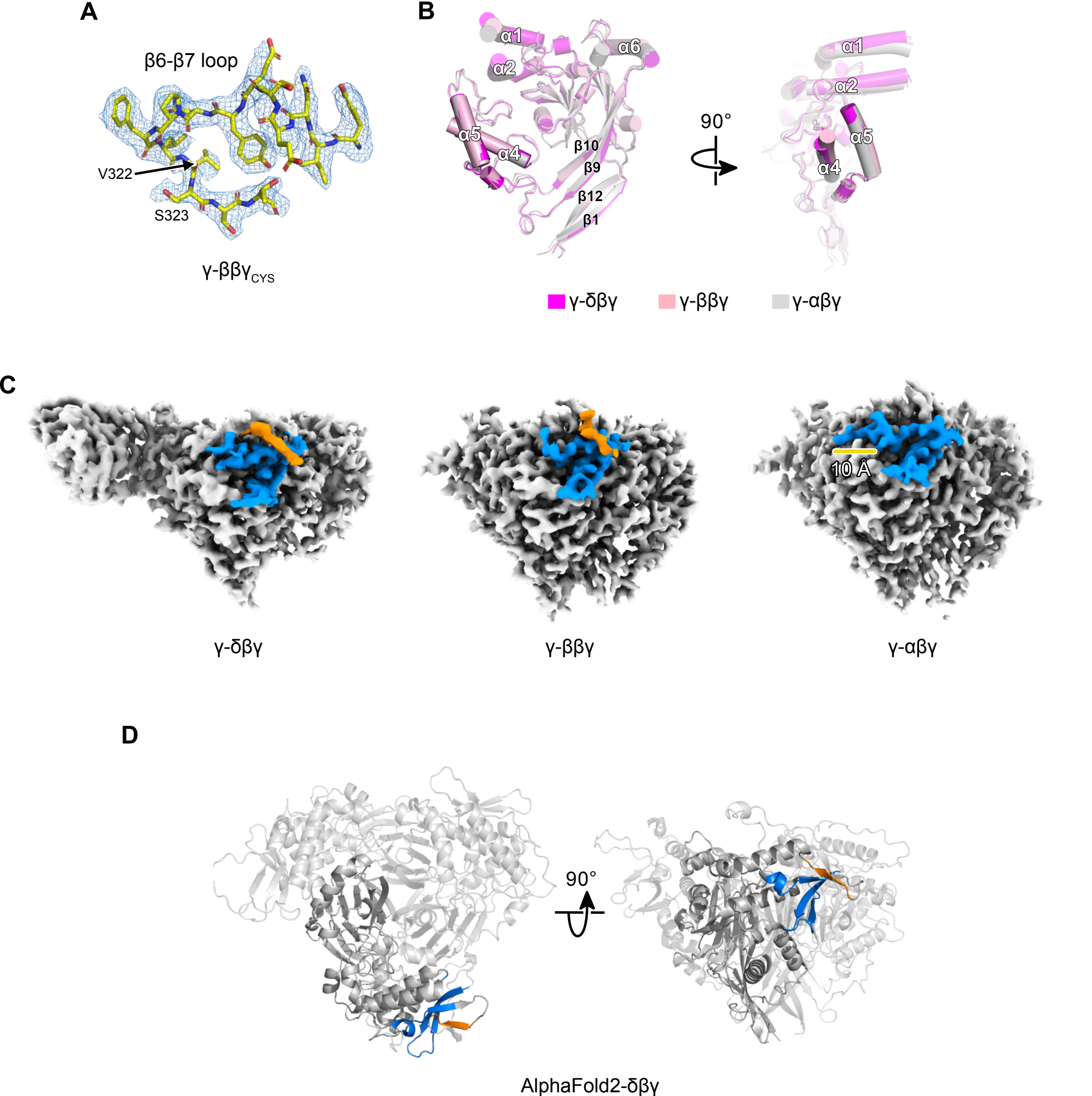

### Figure S6

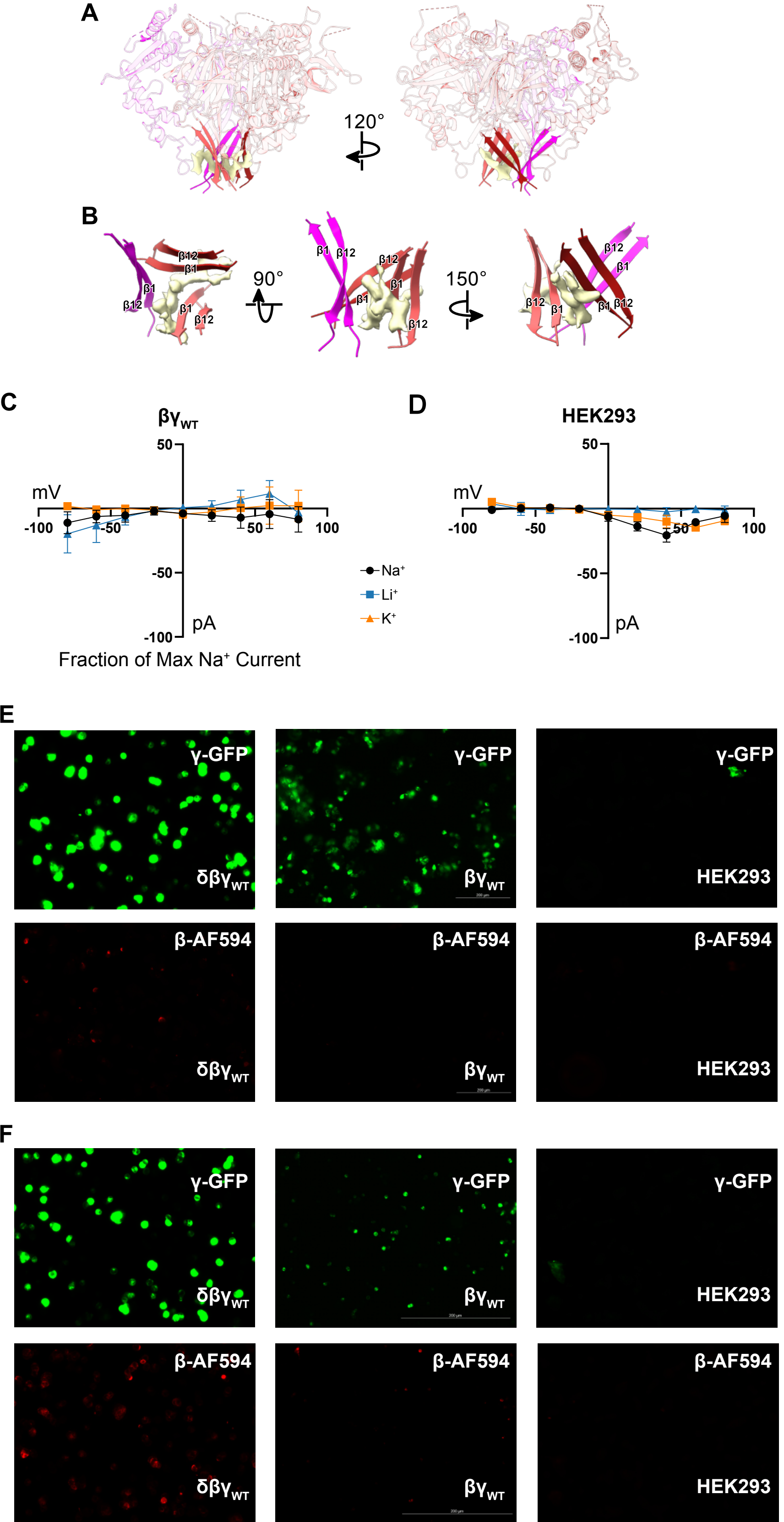

### Figure S7

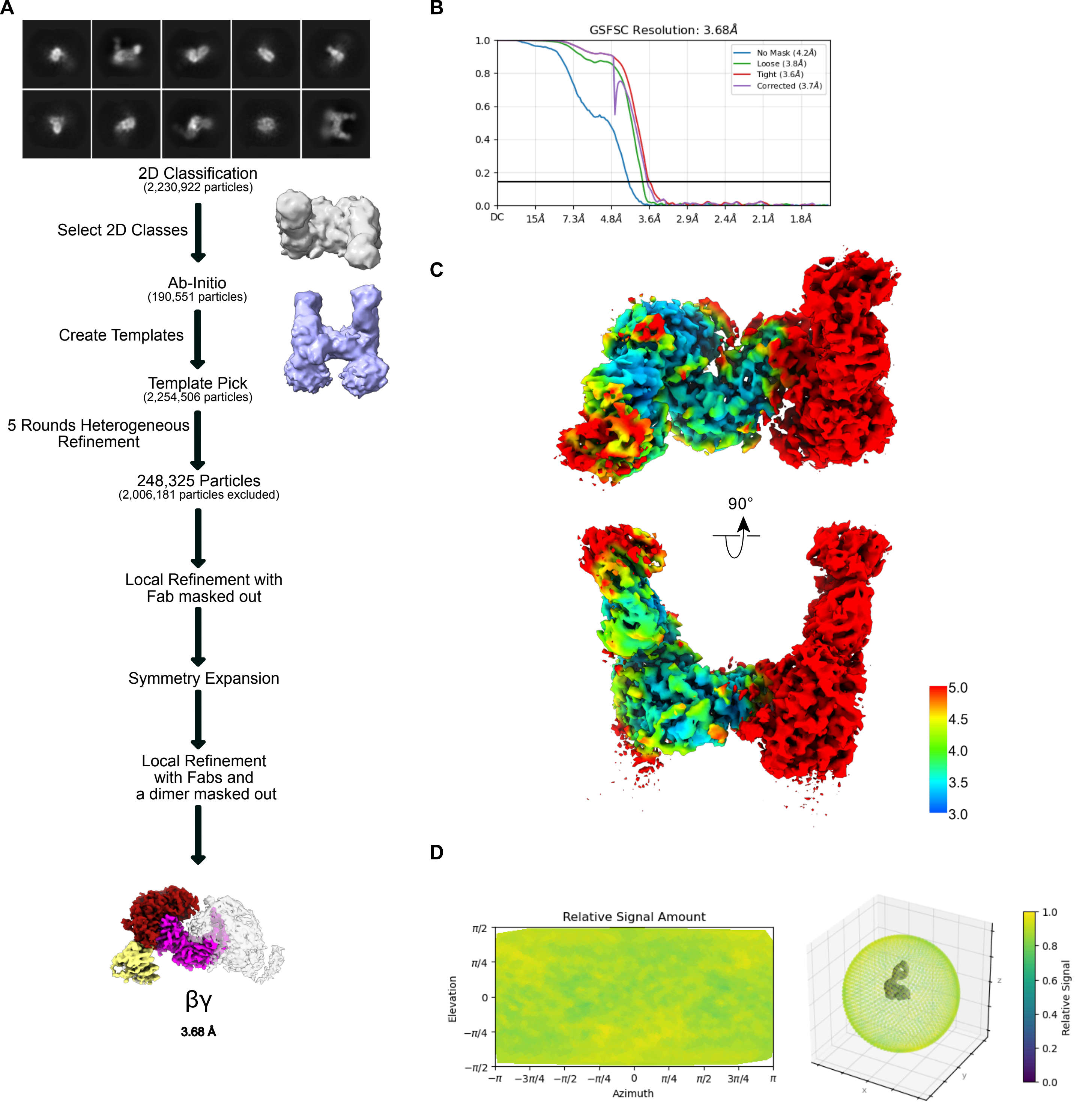
