## Supplementary material for "Structural Insights into Subunit-Dependent Functional Regulation in Epithelial Sodium Channels": Table S1

**Table S1. Cryo-EM data collection, refinement, and validation statistics**

| | δβγ (EMD-44674) (PDB: 9BLR) | ββγ (EMD-44889) (PDB: 9BTG) | βγ-βγ (EMD-44896) (PDB: 9BTU) |
|---|---|---|---|
| Data Collection and processing | | | |
| Magnification (kX) | 165 | 165 | 165 |
| Voltage (kV) | 300 | 300 | 300 |
| Electron exposure (e-/Å$^2$) | 50 | 50 | 50 |
| Defocus range (μm) | -0.8 to -2.5 | -0.8 to -2.5 | -0.8 to -2.5 |
| Raw pixel size (Å) | 0.40075 | 0.40075 | 0.413 |
| Symmetry imposed | C1 | C1 | C1 |
| Initial particle images (no.) | 4,884,175 | 4,884,175 | 1,121,564 |
| Final particle images (no.) | 192,941 | 342,011 | 248,323 |
| Map resolution (Å) | 3.38 | 3.12 | 3.68 |
| FSC threshold | 0.143 | 0.143 | 0.143 |
| Map resolution range (Å) | 3.56 - 3.18 | 3.24 - 2.90 | 3.86 - 3.44 |
| Refinement | | | |
| Initial model used (PDB code) | 6WTH | 6WTH | 6WTH |
| Model resolution (Å) | 3.5 | 3.3 | 3.9 |
| FSC threshold | 0.5 | 0.5 | 0.5 |
| Map sharpening B factor (Å$^2$) | -99 | -101 | -147 |
| Model composition | | | |
| Non-hydrogen atoms | 10,504 | 9,869 | 7,859 |
| Protein residues | 1,169 | 1,207 | 837 |
| Ligands | 14 | 12 | 16 |
| B factors (Å$^2$) | | | |
| Protein | 65.58 | 64.15 | 146.73 |
| Ligand | 110.58 | 83.25 | 135.71 |
| Root mean square deviations | | | |
| Bond lengths (Å) | 0.006 | 0.005 | 0.005 |
| Bond angles (°) | 0.942 | 0.744 | 0.807 |
| Validation | | | |
| MolProbity score | 0.83 | 0.65 | 1.14 |
| Clashscore | 0.45 | 0.41 | 1.63 |
| Poor rotamers (%) | 0.00 | 0.28 | 0.13 |
| Ramachandran plot | | | |
| Favored (%) | 97.04 | 97.97 | 96.48 |
| Allowed (%) | 2.96 | 2.03 | 3.52 |
| Disallowed (%) | 0.00 | 0.00 | 0.00 |
