## Supplementary material for "Structural Insights into Subunit-Dependent Functional Regulation in Epithelial Sodium Channels": Table S2

### Table S2. Root-mean-square deviation of Cα atoms in each subunit domains

| | RMSD (subunit superposition) | RMSD (trimer superposition) | RMSD (trimer superposition with β as reference) |
|---|---|---|---|
| $\delta$-$\delta\beta\gamma_{CYS}$ vs $\alpha$-$\alpha\beta\gamma_{6WTH}$ | 1.0<br>2.4<br>2.2<br>1.4<br>1.7 | 1.1<br>2.3<br>2.6<br>2.1<br>3.5 | 1.8<br>2.4<br>3.3<br>3.1<br>5.1 |
| $\beta$-$\delta\beta\gamma_{CYS}$ vs $\beta$-$\alpha\beta\gamma_{6WTH}$ | 0.4<br>1.2<br>0.4<br>0.4<br>0.5 | 0.8<br>1.0<br>0.8<br>0.6<br>1.0 | 0.4<br>1.2<br>0.4<br>0.4<br>0.5 |
| $\gamma$-$\delta\beta\gamma_{CYS}$ vs $\gamma$-$\alpha\beta\gamma_{6WTH}$ | 0.6<br>0.7<br>0.5<br>1.8<br>0.9 | 0.6<br>0.6<br>0.5<br>1.8<br>1.1 | 0.5<br>0.5<br>0.8<br>2.2<br>1.6 |
| $\delta$-$\delta\beta\gamma_{CYS}$ vs $\beta^1$-$\beta\beta\gamma_{CYS}$ | 0.9<br>1.9<br>2.1<br>1.7<br>3.0 | 0.7<br>2.0<br>2.1<br>1.8<br>3.6 | 0.9<br>2.0<br>2.1<br>2.2<br>3.5 |
| $\beta$-$\delta\beta\gamma_{CYS}$ vs $\beta^2$-$\beta\beta\gamma_{CYS}$ | 0.4<br>1.0<br>0.3<br>0.3<br>0.3 | 0.5<br>1.0<br>0.5<br>0.4<br>0.4 | 0.4<br>1.0<br>0.3<br>0.3<br>0.4 |
| $\gamma$-$\delta\beta\gamma_{CYS}$ vs $\gamma$-$\beta\beta\gamma_{CYS}$ | 0.5<br>0.7<br>0.4<br>1.3<br>0.7 | 0.3<br>0.5<br>0.5<br>1.4<br>0.9 | 0.4<br>0.4<br>0.8<br>1.5<br>1.2 |
| $\beta^1$-$\beta\beta\gamma_{CYS}$ vs $\alpha$-$\alpha\beta\gamma_{6WTH}$ | 0.8<br>1.8<br>1.9<br>1.7<br>2.4 | 1.0<br>1.0<br>2.5<br>2.3<br>5.3 | 1.6<br>1.4<br>3.1<br>3.3<br>6.6 |
| $\beta^2$-$\beta\beta\gamma_{CYS}$ vs $\beta$-$\alpha\beta\gamma_{6WTH}$ | 0.5<br>1.1<br>0.4<br>0.4<br>0.4 | 0.7<br>1.4<br>0.7<br>0.5<br>0.9 | 0.5<br>1.1<br>0.4<br>0.4<br>0.4 |
| $\gamma$-$\beta\beta\gamma_{CYS}$ vs $\gamma$-$\alpha\beta\gamma_{6WTH}$ | 0.4<br>0.9<br>0.6<br>1.1<br>0.8 | 0.5<br>0.4<br>0.6<br>0.9<br>1.4 | 0.3<br>0.4<br>0.7<br>1.23<br>2.1 |

**Upper palm**
**Knuckle**
**β-ball**
**Finger**
**Thumb**
